# Mobile species’ responses to surrounding land use generate trade-offs among nature’s contributions to people

**DOI:** 10.1101/2025.03.11.638535

**Authors:** Sophie A. O’Brien, Jason M. Tylianakis, Dean P. Anderson, Andrea Larissa Boesing, Hao Ran Lai, Gaëtane Le Provost, Peter Manning, Margot Neyret, Nico Blüthgen, Kirsten Jung, Paul Magdon, Sandra Müller, Michael Scherer-Lorenzen, Noëlle Schenk, Sandra Lavorel

## Abstract

Agricultural landscapes provide material, non-material and regulating contributions that affect human wellbeing. The responses of these nature’s contributions to people (NCP) to land-use patterns depend on supporting biota with different habitat requirements, generating trade-offs and synergies. Predictions of NCP trade-offs could inform land-use decisions, but these do not typically account for the effects of land-use patterns on the movement of NCP-providing species, nor for interactions among NCP providers. To explore spatial trade-offs and synergies in eight indicators of NCP, we used Bayesian models that allow for interactions among land uses and among NCP using data from 150 grassland sites across rural Germany. We found that spatial arrangements of forest and open habitat influenced many NCP: acoustic diversity, birdwatching potential, natural enemy abundance and pollination. In particular, the amount and proximity of land uses in the surrounding landscape, especially forest and open habitat, drove the supply of most NCP. However, NCP provided by smaller-bodied taxa (herbivory and pathogen infection) responded weakly to landscape factors. Multiple NCP provided by a given taxon responded differently to their surrounding landscape (e.g. bird-provided caterpillar predation and seed predation), leading to trade-offs among NCP over short distances (≤60 m). These were caused by different rates and directions of response to amount and location of land uses. Resulting spatial predictions revealed that grassland-dominated or mixed forest-grassland areas better maximize beneficial NCP and minimize detrimental NCP across landscapes than purely forest-dominated areas. This suggests promoting semi-natural vegetation in agricultural landscapes to provide greater-than-additive benefits to net NCP supply.

**Significance Statement:** Land use affects nature’s contributions to people (NCP), including those provided by mobile species, in complex ways. Variation in the responses of species to the amount and location of land uses results in NCP trade-offs across landscapes, but decision-makers lack the capacity to spatially predict these trade-offs. We predict how the supply of both beneficial and detrimental NCP and associated trade-offs vary across diverse rural landscapes and find that grassland-dominated or mixed forest-grassland areas deliver higher net NCP supply than forest-dominated areas in our study system. Our findings support policies for promoting semi-natural vegetation cover in agricultural landscapes, as these may provide non-additive benefits for NCP.

## Main Text

Global change alters the ability of landscapes to supply nature’s contributions to people (NCP), “all the contributions, both positive and negative, of living nature to people’s quality of life” (Díaz et al. 2018). Globally, two-thirds of human-modified landscapes lack the natural and semi-natural habitat required for the supply of multiple NCP (Mohamed et al. 2024). In efforts to reverse this global trend, many countries and regions are adopting policies for increasing semi-natural vegetation cover and decreasing land-use intensity (Convention on Biological Diversity 2022). However, depending on its intensity, land use has contrasting influences on co-occurring NCP (NCP bundles), such as pollination and crop production (Neyret et al. 2021, Simons et al. 2021). This generates trade-offs and synergies among NCP in mixed-use, mixed-intensity landscapes (Le Provost et al. 2023). Moreover, because landscape processes influence the capacity of natural areas to provide NCP (Garibaldi et al. 2011), predictions of the provision of NCP bundles for use in land-use decisions must account for landscape context (Metzger et al. 2021). Yet, three outstanding gaps limit such predictions.

Firstly, many NCP are provided by particular species, whose presence and abundance is determined by a combination of local- and landscape-level abiotic and biotic environmental conditions (Kass et al. 2024). It is therefore challenging to predict spatial variation in NCP trade-offs and synergies when species movement across land uses or lateral physical processes determine NCP flow across landscapes (Blitzer et al. 2012, Han et al. 2024, O’Brien et al. 2024). Substantial work has demonstrated how species-provided NCP respond to surrounding landscape composition (Lander et al. 2011, Stein et al. 2014, Hohlenwerger et al. 2024), but such approaches do not account for the spatial arrangement of multiple habitats and therefore cannot predict well into new landscapes. Analyzing how species and associated NCP change across the edges of a focal land-use (Ricketts et al. 2008) can capture distance decay of NCP from a source habitat. However, such designs do not account for multiple patches of multiple land uses, which can act as both sources and sinks of NCP-providing organisms. Moreover, the ways in which different NCP are affected by the surrounding land uses may also depend on the taxa that contribute to each NCP. For example, larger taxa (e.g. birds) may disperse further than smaller taxa (e.g. arthropods; (Jenkins et al. 2007)). Conversely, larger-bodied species are often more sensitive to land-use intensity and change (Newbold et al. 2013), and the NCP they provide could therefore exhibit steeper distance decay from preferred habitat.

Secondly, a range of mechanisms may cause interactions between the effects of different land uses on NCP supply. For example, species may use complementary resources from multiple land uses (Stein et al. 2014) or adjacent land uses may distract mobile species, such as when pollinating arthropods or mammals are distracted by neighboring resource-rich patches (Lander et al. 2011). Drawing from movement ecology and landscape ecology, the movement of mobile species is dependent on both their characteristics (e.g. potential mobility, degree of generalism and feeding guild; (Blitzer et al. 2012, Peña et al. 2023)) and their fine-scale movement decisions within increasingly fragmented and modified landscapes (Seidel et al. 2018). Thus, trade-offs among NCP arise from differing tendencies of different NCP-providing organisms to move among land uses of varying distance.

Thirdly, there are trade-offs among beneficial and detrimental NCP (Díaz et al. 2018). Even a given taxon can concurrently provide beneficial and detrimental NCP, for example small rodents consume weed seeds but also damage crops (Fischer et al. 2018). Yet, our ability to predict the net supply of NCP at a given point in space (netNCP; (Neyret et al. 2024)) is presently limited because most previous work does not integrate detrimental NCP (Spake et al. 2017, Simons et al. 2021).

We fill these three gaps with a novel, spatially-explicit approach for predicting NCP trade-offs that accounts for the amount, location, and identity of land-use patches across landscapes, whilst allowing for interactive effects between land uses and between different NCP. We explore how the supply of five beneficial (pollination, natural enemy abundance, birdwatching potential, caterpillar predation and acoustic diversity) and three detrimental (seed predation, herbivory and pathogen infection) NCP indicators (hereafter, ‘NCP’), provided in grasslands by mobile species, vary spatially with the distribution of land uses in the surrounding landscape, across three European agricultural regions. We test four hypotheses: (H_1_) Different NCP are favored by different land uses, and NCP that share a providing species will respond similarly to similar landscape patterns (e.g. all bird-provided NCP will respond to the availability of the same land uses; Fig. 1; *SI Appendix* Table S1); (H_2_) the combined effect of multiple land uses on NCP provided by mobile species will be non-additive, because adjacent land uses may distract mobile species (Lander et al. 2011) or because species with a broad diet may benefit from the resources provided by multiple land uses (Stein et al. 2014) (e.g. merged arrows to arthropod-provided pollination in Figure 1); (H_3_) the supply of multiple NCP will vary spatially, generating trade-offs caused by differences in the direction and/or magnitude of NCP responses to different land uses (and their arrangement); and (H_4_) NCP provided by larger-bodied taxa, such as birds, will exhibit flatter spatial decay from a source land use due to their larger movement ranges.

**Figure 1.**
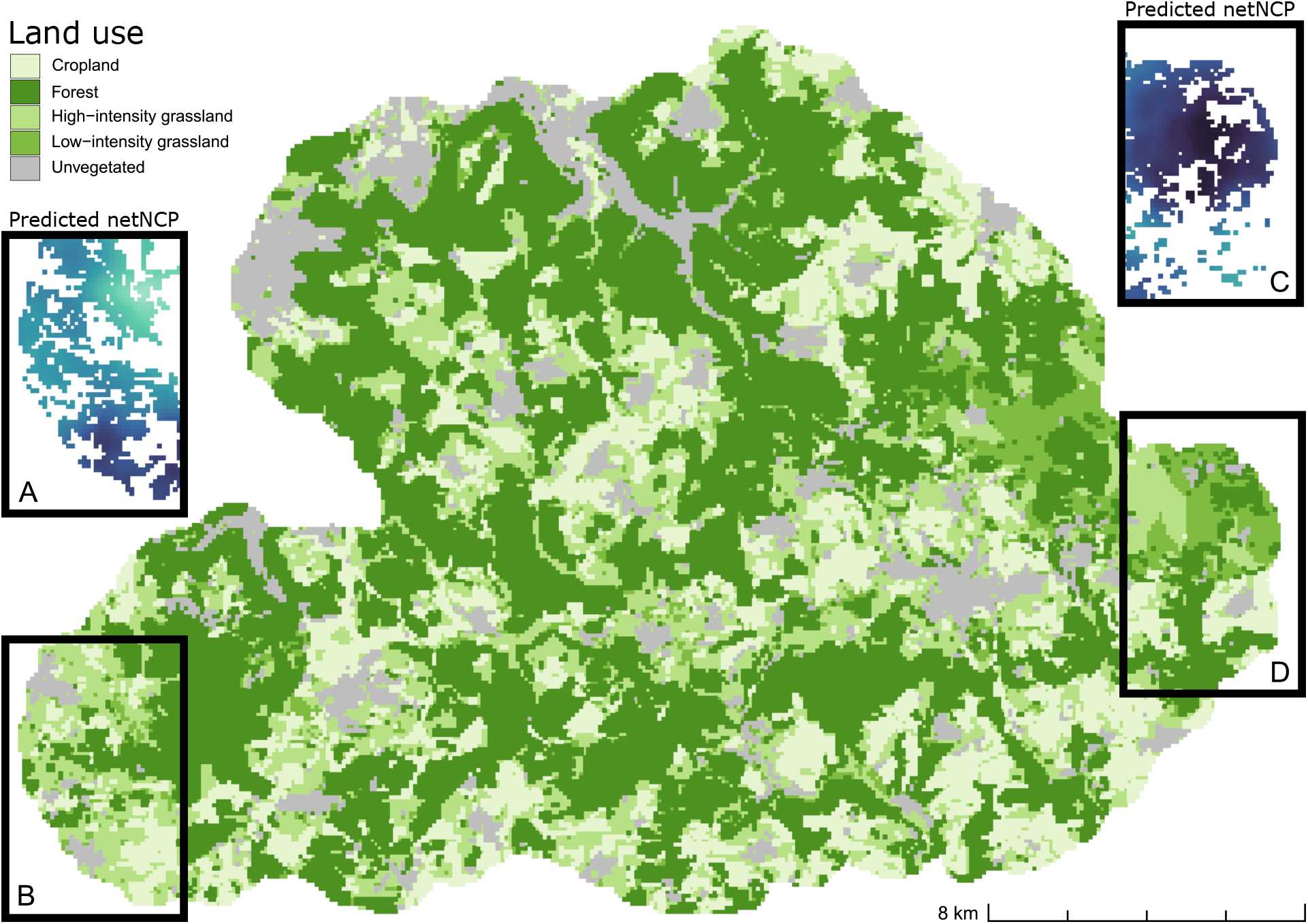
Conceptual summary of the NCP indicators modeled. Dark grey arrows show the land uses in the surrounding landscape expected to be the strongest positive drivers of each NCP (based on existing literature; *SI Appendix* Table S1) in a measured grassland plot. Light grey curved arrows show the hypothesized directional effects between NCP. We expect both low-intensity grassland and forest to strongly drive NCP supply of pollination (merging dark grey arrow), and we test whether these effects are additive or interactive in model selection (see ‘Statistical analysis and model selection’ in Materials and Methods). The mobile providers of each NCP are represented as icons in NCP boxes, where the larger left-most icon represents the primary provider of that NCP if multiple providers are shown. The NCP deemed beneficial are presented in blue boxes, and those deemed detrimental in red boxes. The indicators measured for each NCP are as follows: natural enemy abundance (number of brood cells recorded in trap nests attacked by parasitoids of pest arthropods); pollination (number of flower visitors); acoustic diversity (the distribution of acoustic energy among frequency bands during diurnal recordings); birdwatching potential (bird species richness); caterpillar predation (probability of dummy caterpillar predation by birds after 48 hours); pathogen infection (total cover of foliar fungal pathogens); seed predation (probability of sunflower seed removal after 48 hours); and herbivory (total proportion of leaf area damaged by invertebrate herbivores).

We test these hypotheses using comprehensive NCP and land-use datasets for 150 agricultural grassland plots of the German Biodiversity Exploratories project (Fischer et al. 2010). We extend existing spatial modelling techniques (Clark et al. 1999, Ricketts et al. 2008, O’Brien et al. 2024) to illustrate how point-based measures of multiple NCP can be used to generate a high-resolution prediction surface of netNCP supply that accounts for the direct and indirect effects (via another NCP; Fig. 1 and *SI Appendix* Table S1) of multiple land uses. We apply a stepwise model selection approach to determine the relative importance of the distance-weighted amount of four land uses (forest, cropland, low-intensity grassland, and high-intensity grassland), all two-way land-use interactions, environmental covariates, and other NCP. Across all models, we fit NCP-specific distance-decay parameters to test for between-NCP differences in the spatial range of influence. The eight NCP, considered as beneficial or detrimental in these European agricultural landscapes, and their hypothesized responses to land use and to other NCP are summarized in Figure 1.

## Results

### Differing responses of NCP to the amount and proximity of land uses generate trade-offs

Overall, the supply of most NCP in grasslands was driven to some extent by the amount and proximity of different land use types in the surrounding landscape, but differences in specific NCP responses generated trade-offs (Fig. 2, *SI Appendix* Table S2). All four land-use main effects and two-way interactions were removed in model selection except for those described below and presented in Figure 2 (see ‘Statistical analysis and model selection’ in Materials and Methods). The control variable *TWI*_*i*_ (topographic wetness index) was retained in the best-fitting models for birdwatching potential, natural enemy abundance, pollination, and seed predation; secondary NCP variables were retained in the best-fitting models for pathogen infection (herbivory) and caterpillar predation (birdwatching potential; *SI Appendix* Table S2).

**Figure 2.**
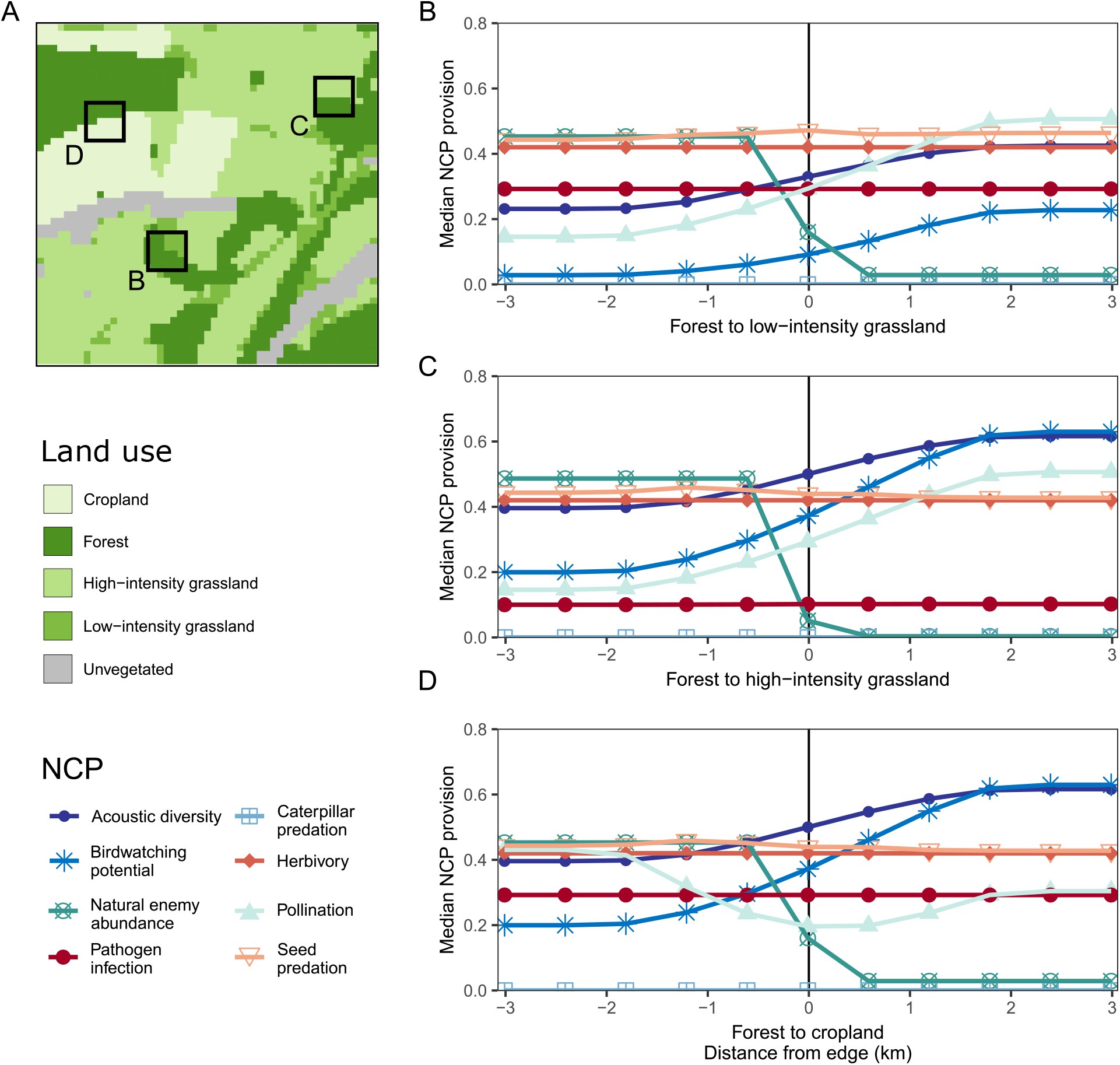
Parameter estimates (points and error bars) and variance explained (vertical bars) from the best-fit models of each NCP, separated by region. Parameters for forest, cropland, low- and high-intensity grassland are shown in green, orange, yellow and dark yellow, respectively. Purple points and bars correspond to topographic wetness index (TWI; which has shown to be an important predictor of multiple NCP in the region), and blue points and bars to NCP covariates when retained in the best-fit model of the focal NCP. All two-way land-use effect interactions are shown in brown. Points and error bars are posterior median and 90% credible intervals, respectively. Credible intervals around the coefficient estimates that overlap with zero indicate that the strength of the effect is weak, although the variable’s retention in model selection suggests their inclusion did improve model fit. Vertical bars to the right of each region column shows the NCP-specific proportions of explained variance attributable to each predictor, by region. The NCP herbivory is omitted from this figure because the best-fit model retained only the intercept term (not presented here).

Of the land uses we tested, the amount of forest surrounding grassland was a prevalent driver retained in model selection for most NCP (Fig. 2, *SI Appendix* Table S2). These results were robust to the scale at which land-use main effects were calculated (*SI Appendix* Figure S1). However, the strength and direction of forest effects varied between NCP. For example, birdwatching potential was strongly negatively associated with high forest area in the surrounding landscape in all regions, while natural enemy abundance responded positively to forest cover in regions with large forest patches (Alb and Hainich; Fig. 3B and *SI Appendix* Table S3).

**Figure 3.**
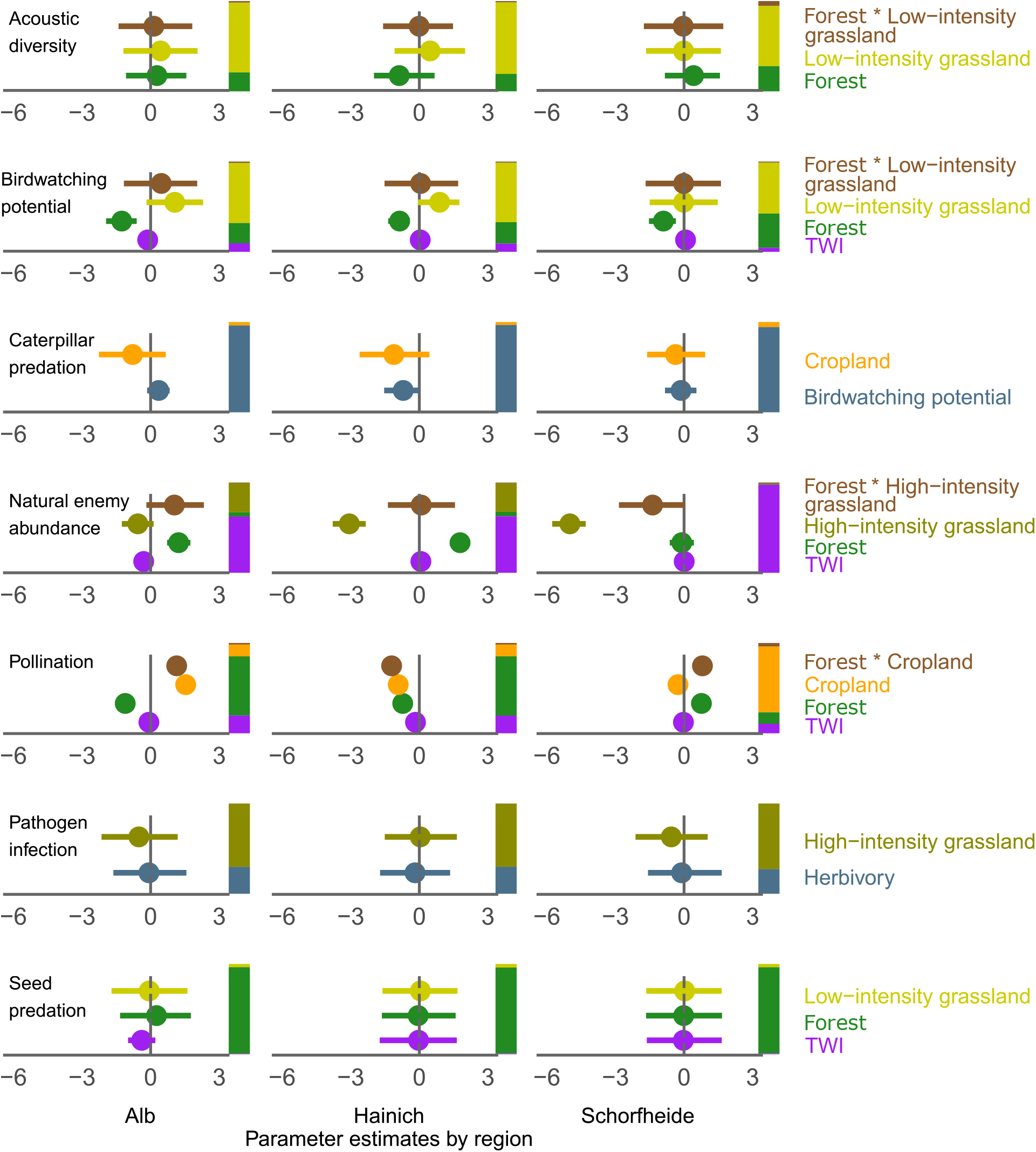
Predicted supply of eight NCP, displayed for a subset of the Schwäbische Alb region (Panel A). Panels B–D show fitted values for the normalized median supply of each NCP in a grassland plot surrounded by different landscape contexts. Landscape contexts represent moving from a landscape of pure forest (negative x-axis values) across the boundary into pure (B) low-intensity grassland; (C) high-intensity grassland and (D) cropland, predicted using the best-fitting models for each NCP in *SI Appendix* Table S2 and coefficient estimates from the Schwäbische Alb region. These between-land-use boundaries correspond to the areas marked B, C and D in Panel A. Spatial base data: © GeoBasis-DE / BKG (2018); Terms of use: http://sg.geodatenzentrum.de/web_public/nutzungsbedingungen.pdf.

NCP provided primarily by birds, which we hypothesized would exhibit flatter spatial decay from a source land use than arthropods (large fitted values of γ in Eq. 1; (Jenkins et al. 2007)), in fact showed the strongest and most consistent responses to the amount and location of land uses in the surrounding landscape. Specifically, bird-provided NCP responded to forest and low-intensity grassland, both directly (acoustic diversity, birdwatching potential and seed predation) and indirectly via another NCP (caterpillar predation via birdwatching potential; Fig. 2 and *SI Appendix* Table S3). Amount and location of low-intensity grassland was more important for acoustic diversity and birdwatching potential, with positive effects in most regions. In contrast, most variation in seed predation was explained by the amount of forest surrounding the grasslands, with lower supply in more forested areas in most regions (Fig. 2 and *SI Appendix* Tables S2 and S3).

Strongly opposing responses of pollination and natural enemy abundance to the surrounding landscape caused trade-offs among NCP provided by arthropods. For example, a grassland with greater surrounding forest area supported significantly higher natural enemy abundance but significantly lower pollination supply (Fig. 2, *SI Appendix* Table S2). In contrast, pollination was more strongly driven by surrounding cropland area, though the direction of this effect differed across regions (Fig. 2, *SI Appendix* Tables S2 and S3). These differences could be attributable to regional differences in landscape composition; i.e. the lowest overall forest cover and smallest forest patch size in the Schorfheide region aligned with a relatively high importance of cropland (Fig. 2B) (Boesing et al. 2024).

Even when multiple NCP responded in the same direction to a given land use, differing strengths of their response generated trade-offs by changing the relative balance of NCP at different distances from source patches. The supply of most NCP decayed steeply with distance from a source land-use patch, indicating that local flows of NCP-providing species are key to supply. Conversely, pollination showed a flatter decay (*SI Appendix* Table S3). This generated trade-offs, for example, because increasing amounts of surrounding forest area in Schorfheide grasslands rapidly maximized acoustic diversity, while more was needed to maximize pollination (mean γ of 2.1 and 20.2 respectively; *SI Appendix* Table S3). Contrary to expectations, NCP provided by birds (which are larger-bodied) were not influenced by land use at greater distances than NCP provided by arthropods. That is, we found no evidence of taxa-specific differences in the distance decay of NCP supply (*SI Appendix* Table S3).

### Effects of land-use interactions on NCP supply

We detected interactions between neighboring land uses for some NCP, although these only accounted for a small proportion of explained variance (Fig. 2B, *SI Appendix* Tables S3). For example, surrounding forest more strongly affected birdwatching potential and acoustic diversity if the landscape also contained grassland managed at high-rather than low-intensity (Fig. 3B).

### Predictability among regions varies across NCP

The responses of bird-provided NCP to the surrounding landscape were consistent across regions (acoustic diversity, birdwatching potential and seed predation; Fig. 2 and *SI Appendix* Table S3). In contrast, NCP provided mostly by arthropods (herbivory, pollination, natural enemy abundance) responded less consistently (Fig. 2). This is represented by among-region variation in the direction and magnitude of the predictor variables in their best-fit models, which likely reflects different landscape contexts (*SI Appendix* Table S3). However, the NCP-NCP interactions captured by covariates in the best-fit models (birdwatching potential in the caterpillar predation model and herbivory in the pathogen infection model) were consistent across regions (Fig. 2). Overall, these findings suggest that both bird-provided NCP and NCP-NCP relationships are generalizable, while those provided by arthropods are less predictable and more specific to regional context.

### Different directions and magnitudes of land-use responses between NCP create trade-offs across landscapes

Accounting for the direct effects of land use on NCP supply and the interactions between land uses and between NCP allowed us to predict individual NCP supply within each study region. We combine these surfaces of individual predicted NCP to predict a landscape-wide surface of netNCP (Fig. 4A and C). As hypothesized, we find that the differing directions and magnitudes of NCP responses to different surrounding land uses (Fig. 2) result in spatial variation in the supply of multiple NCP. In the Schwäbische Alb region (Fig. 4), areas of high net beneficial NCP supply align with areas of grassland and mixed habitats, rather than forest-dominated areas (Figs 4A vs. 4C). We also find that landscape areas with large patches of several land uses, rather than a greater number of smaller patches, support the highest netNCP (Figs 4A vs. 4C). This could be because beneficial NCP showed stronger and more complex responses to surrounding land use than did detrimental NCP (*SI Appendix* Table S2). Although we are unable to state whether this finding is generalizable beyond our study system, these differences do show that there can be stark trade-offs between the landscape structures that promote beneficial and detrimental NCP.

**Figure 4.**
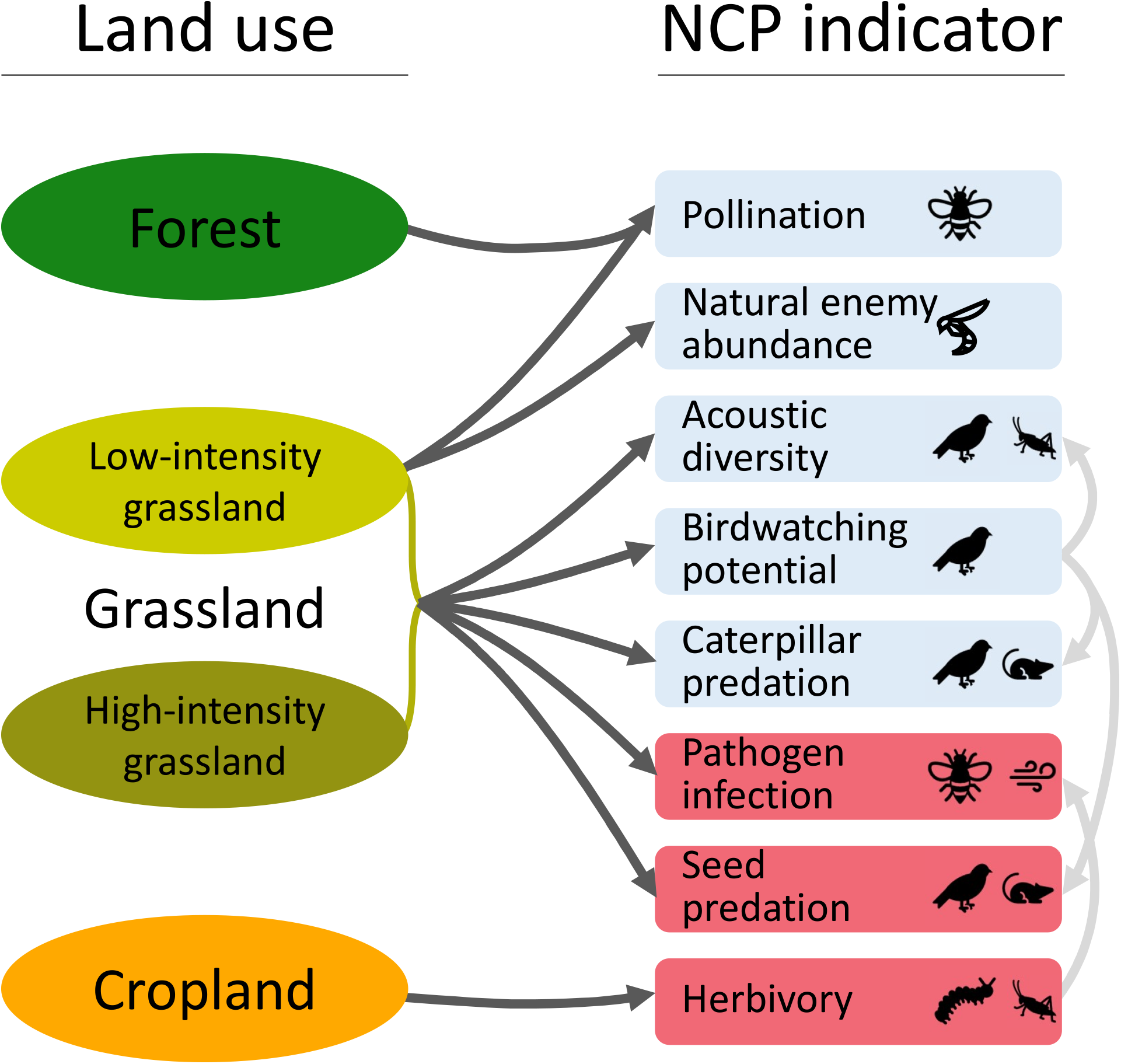
Land use across the Schwäbische Alb region, with heatmaps of median predicted netNCP supply (insets A and C) in grasslands given the surrounding landscape for two subsets of the region (insets B and D). Heat maps in insets A and C correspond to the land-use configurations in insets B and D respectively. Here, darker blue areas represent higher values of netNCP, resulting from relatively higher supply of beneficial NCP or relatively lower supply of detrimental NCP relative to pixels in paler areas. Unfilled pixels in insets A and C correspond to pixels without grassland cover (the land use for which projections were made). Each NCP was predicted individually for each 100 m pixel using best-fitting models from *SI Appendix* Table S2, accounting for direct and indirect effects of habitat (i.e. predicted pathogen infection accounts for indirect land-use effects via predicted herbivory supply, and predicted caterpillar predation via predicted birdwatching potential; *SI Appendix* Table S3). NCP were weighted such that beneficial and detrimental NCP were equally weighted by dividing by the total number of beneficial (five) or detrimental (three) NCP respectively. Spatial base data: © GeoBasis-DE / BKG (2018); Terms of use: http://sg.geodatenzentrum.de/web_public/nutzungsbedingungen.pdf.

## Discussion

Extensive research has demonstrated how NCP supply at a given location is affected by surrounding landscape composition (Ricketts et al. 2008, Le Provost et al. 2023, Hohlenwerger et al. 2024). However, NCP trade-off analysis and prediction urgently need to move beyond spatial association and correlation to incorporate a more mechanistic understanding of drivers (Gomes et al. 2021). Specifically, understanding how the biology of NCP-providing species underpins causal among-NCP and NCP-land use relationships could better inform land management (Dade et al. 2019).

We drew upon seminal methods for estimating dispersal shadows (Clark et al. 1999) and recent advances in modelling mobile NCP provider responses to land use (O’Brien et al. 2024) to generate high-resolution spatial predictions of NCP trade-offs that more accurately account for the spatial dynamics of multiple providers and for complex interactive effects of land uses. Importantly, we explored which trade-off predictions were transferable across landscapes based on generalities and context dependencies we identified; the absence of such validation is a persistent weakness of many approaches (Spake et al. 2017, Kass et al. 2024). Our results reveal that the amount and identity of land uses in the landscape surrounding grasslands drive the supply of most NCP, both directly and indirectly (via another NCP). Importantly, the NCP interaction effects we found are driven by the biology of NCP-providing taxa, thereby moving beyond broad scale patterns of correlation or co-occurrence (Dade et al. 2019, Gomes et al. 2021). Additionally we revealed interactions among land uses, with the responses of most mobile NCP-providing taxa to a given land use being modified by neighboring land use types.

Contrary to expectations, NCP provided by the same taxa did not respond consistently to their surrounding landscape. For example, some NCP associated with bird diversity (acoustic diversity and birdwatching potential) consistently responded positively to the area of open land uses in all regions. This finding aligns with the general pattern of European bird diversity being better supported by mixed and semi-open habitats than extensive forest (Wehner et al. 2020). Conversely, other NCP provided by birds through trophic interactions (caterpillar predation and seed predation) were insensitive to surrounding land use (Figs. 2 and 3B–D), possibly because many species which feed heavily upon caterpillars are forest-dwelling (e.g. *Parus major*) (Cholewa and Wesołowski 2011). Favorable land uses for arthropod-provided NCP also varied considerably (Figs. 2 and 3B–D). These inconsistencies align with trait differences among taxa that contribute to different ecosystem functions and NCP. For example, although birdwatching potential, caterpillar predation and seed predation are all provided by birds, it is known that birds with different diets vary in their dispersal traits (Peña et al. 2023). Similarly, arthropods in higher trophic levels tend to be more generalist and therefore more likely to disperse into nearby habitats (Blitzer et al. 2012). This may explain why surrounding land uses played a more important role for natural enemy abundance than for herbivory or pathogen infection.

When we categorize the indicators of our focal NCP as either diversity-driven (acoustic diversity, birdwatching potential, natural enemy abundance, pollination) or interaction-driven (seed predation, caterpillar predation, herbivory and pathogen infection), interaction-driven NCP consistently show weaker responses to land use in the surrounding landscape (Fig. 3B–D). In contrast, inclusion of candidate mechanistic linkages among NCP improved model fit for interaction-driven (caterpillar predation and pathogen infection) but not diversity-driven (acoustic diversity) NCP. This supports recent calls for building species interactions into NCP trade-off predictions to improve accuracy and realism (Kass et al. 2024), especially when the indicators used as proxies for NCP measure interactions, not diversity.

Surprisingly, we did not find the hypothesized taxonomic differences in the distance decay among the NCP tested. We expected larger-bodied taxa to have the flattest distance decay, and NCP involving specialist interactions to be restricted to a single land use and thus show steep distance decay (Blitzer et al. 2012). Instead, we found consistently steep distance decay for all NCP except pollination. The flatter distance decay of pollination means that its supply responds to land-use changes over a wider area than the other NCP. Such differences in distance decays create spatial variability in trade-offs. The lack of taxa-specific differences echoes a recent finding that, against expectations based on body size and movement range, bird-provided NCP were not affected at broader scales than arthropod-provided NCP (Hohlenwerger et al. 2024). Overall, the lack of common response of NCP provided by the same taxa to land use suggests that management decisions about NCP need to consider the spatial processes underlying specific NCP (Rey et al. 2024).

We tested the extent to which interactions among adjacent land uses result in interactions among NCP. Here, trade-offs were generated by differing NCP responses to surrounding landscape composition (Fig. 3). Trade-offs can emerge if a land-use change increases the supply of one beneficial NCP whilst reducing another, for example where increasing the forest area surrounding a grassland increases natural enemy abundance but decreases pollination (Figs. 3B–D). However, trade-offs can also emerge when a beneficial and a detrimental NCP respond in the same direction to a land-use type. For example, seed predation and natural enemy abundance both responded positively to surrounding forest area (Fig. 3D). The different slopes of those responses will change the balance of that trade-off as surrounding forest increases. Moreover, we find that initial increases in surrounding forest cover will rapidly increase natural enemy abundance more than seed predation, but further increases will keep increasing seed predation relative to a stabilized level of natural enemy abundance (Fig. 3C). We therefore add to previous studies quantifying trade-offs among beneficial NCP (Spake et al. 2017, Le Provost et al. 2023) by incorporating a range of beneficial and detrimental NCP into our high-resolution spatial predictions of the surface of spatial NCP trade-offs.

Our findings are subject to several caveats. Firstly, our approach summed the area of land use weighted by each pixel’s distance from a focal point, but did not account for land-use patch shape or the potential for connectivity and edge effects (Vanneste et al. 2024). Accounting for this additional nuance could improve model predictions, but could come at the expense of generalizability. Secondly, future work could also consider more specific land-use categories than those used in our study, which could allow comparisons of forests of different types and management intensities. Thirdly, our approach was neutral to stakeholder preferences; we assigned weightings to NCP such that total beneficial NCP and total detrimental NCP were equally weighted when visualizing trade-offs (Fig. 4). However, the perception of a NCP as beneficial or detrimental, and their relative importance, depends on a given stakeholder’s cultural and socioeconomic context (Díaz et al. 2018). Our approach would be strengthened in future work by weighting NCP according to spatial preferences, for example using spatial multi-criteria decision analysis (Adem Esmail and Geneletti 2018), as well as incorporating the level of uncertainty around netNCP estimates. Fourthly, our findings of the land-use arrangements that maximize particular NCP (and netNCP) should be interpreted with caution, in part because the predictability of NCP supply (i.e. the consistency of responses across our three study regions) varied among NCP, especially among those provided by arthropods. Nevertheless, our results demonstrate the potential of our approach to provide insights into how multiple NCP co-vary across diverse landscapes, and the factors that drive this covariation.

Land-use change and its interactions with other global change drivers necessitate predictive mapping of NCP provided by mobile species (Le Provost et al. 2023, Han et al. 2024). Accordingly, we present several recommendations based on our findings. Firstly, our finding that the beneficial NCP varied more within landscapes than detrimental NCP suggests that the net balance of NCP supply will be highest close to hotspots of beneficial NCP providers, and thus with a landscape composition that promotes these biota. This finding could be used to guide management that seeks to maximize netNCP (Spake et al. 2017). Secondly, the lack of taxa-specific commonalities in land-use responses indicates that managing bundles of NCP requires more in-depth consideration of the functional mechanisms underlying NCP responses to land use. Thirdly, we suggest that the relevance of diversity-based indicators for NCP that are underpinned by species interactions (e.g. the plant-arthropod mutualism of pollination) needs further scrutiny. More broadly, our findings demonstrate that promoting semi-natural vegetation in agricultural landscapes may provide greater-than-additive benefits to net NCP supply, and management must carefully consider the differing responses of multiple NCP to land-use changes to successfully foster the multifunctionality upon which rural communities rely (Hohlenwerger et al. 2024).

## Materials and Methods

### Study regions and land cover data

We studied 150 agricultural grassland plots (50 × 50 m) evenly divided among three regions of Germany: Schwäbische Alb (Alb), Hainich-Dün (Hainich) and Schorfheide-Chorin (Schorfheide). These regions vary by land cover, soil type and topography, and the plots are part of a long-term research project Biodiversity Exploratories ((Fischer et al. 2010); www.biodiversity-exploratories.de). All plots had been grassland for at least 10 years prior to project commencement. Of the regions, Alb has greatest grassland area (Alb: 35.8%; Hainich: 17.6%; Schorfheide: 17.1%), Hainich the highest proportion of croplands (Alb: 17.2%; Hainich: 52.1%; Schorfheide: 23.6%), and Schorfheide the greatest forest area (Alb: 44.0%; Hainich: 26.3%; Schorfheide: 50.9%) (*SI Appendix* Table S4). The regions also varied in spatial configuration: Alb has the highest density of forest and grassland edges, Hainich the least fragmented landscape overall, and Schorfheide the smallest mean patch size for forests but the largest for grasslands when considering only the main highways crossing them (Boesing et al. 2024).

To proxy for land use, we used land cover data from the Federal Agency of Cartography and Geodesy (Magdon 2023), grouped into four broad categories: forest (including forest and scrub), grassland, cropland, and unvegetated (including water bodies, roads, urban areas and rock) (*SI Appendix* Table S4). Unvegetated areas were not considered in our analyses as these act as barriers to, rather than sources of, NCP. Because management type is known to have considerable effects on NCP supply in grasslands (Neyret et al. 2021), we further classified grassland as either low-intensity or intermediate/high-intensity (hereafter, high-intensity) using a secondary remote-sensed land-use intensity (LUI) dataset that takes into account grazing intensity, mowing frequency and fertilizer application (Lange et al. 2022). We classified each 20 m grid cell of grassland for the whole region as high intensity if the LUI value > 1.2, or low intensity if LUI value ≤ 1.2 (4). For the 8% of land that was classified as grassland according to our primary dataset but not considered grassland in our secondary dataset, we took the LUI value from the nearest grassland pixel with a LUI value.

### Landscape explanatory variables

We first constructed a land-use effect (*LUE*_*l,i*_) variable for cropland, low-intensity grassland, high-intensity grassland and forest. These variables captured the distance-weighted contribution of each focal land use, accounting for the area of a land use *l* and its proximity *d* to a given plot *i*. To calculate *LUE*_*l,i*_ for each plot, we first gridded the landscapes and assigned to each 20 × 20 m cell the majority land use within. We positioned plots in this landscape based on their GPS coordinates. Because species respond at different scales, for each focal NCP we calculated the *LUE*_*l,i*_ variables at the scale previously demonstrated to best explain the response of the primary trophic group that provides that focal NCP, either 500, 1000 or 2000 m; *SI Appendix* Table S5). Here, we constructed a circular buffer with a radius around each plot according to that response scale described in *SI Appendix* Table S5. For each plot, we measured the distance to all cell centroids that fell within its buffer (median number of cells: 31,414 (2000 m), 7,854 (1000 m), 1,962 (500 m)). We modelled the potential effect of each cell of each land use within this buffer, weighted by their distance from a plot. Then we calculated *LUE*_*l,i*_ as:

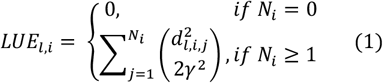

where 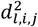 is the Euclidean distance from focal plot *i* to the centroid of cell *j* of land use *l, γ* is a distance-decay parameter that reduced the effect of cell *j* with increasing distance to plot *i*, and *N*_*i*_ is the total number of plots of land use *l* surrounding plot *i*. We used a half-normal distance-decay function to allow for a cell’s effect to diminish with increasing distance from a plot (eq. 1). One *γ* was estimated for all land uses for each NCP to reduce parameterization and because we assumed that species’ mobility is equal across land-use types (although see (Silveira et al. 2016)).

In addition to the *LUE*_*l,i*_ variables and the two-way interactions among them, we controlled for study region (a categorical variable with three levels) and the topographic wetness index (Manning 2023) of plot *i*, an index that accounts for relevant spatial hydrological processes and was demonstrated to be an important factor influencing species in other studies of these grasslands (Le Provost et al. 2023).

### NCP data

We follow Le Provost et al. (2023) and explored the six cultural and aboveground regulating indicators of NCP (‘services and disservices’ in their study, ‘NCP’ hereafter) that are delivered by mobile species. We also included data for two additional NCP delivered by mobile species in the same grassland plots. All NCP were measured and data collected at the plot level (*SI Appendix* Table S6), therefore covering a subset of the study regions’ total environments. We assigned each NCP as either beneficial or detrimental in the context of the German agricultural landscapes in which they were measured, acknowledging that this assignment is stakeholder- and context-specific ((O’Brien et al. 2024); *SI Appendix* Table S1). Overall, our analyses included five beneficial NCP: acoustic diversity (the distribution of acoustic energy among frequency bands during diurnal recordings); birdwatching potential (bird species richness); natural enemy abundance (number of brood cells recorded in trap nests attacked by parasitoids of pest arthropods); pollination (number of flower visitors); and caterpillar predation (probability of dummy caterpillar predation by birds after 48 hours). We also included three detrimental NCP: pathogen infection (total cover of foliar fungal pathogens); herbivory (total proportion of leaf area damaged by invertebrate herbivores); and seed predation (probability of sunflower seed removal after 48 hours). Further details of sampling periods and methods are provided in *SI Appendix* Table S6.

Recent reviews of NCP prediction have called for better integration of interactions among species that provide NCP (Kass et al. 2024). Therefore, we explored ecologically mechanistic potential linkages among NCP according to existing literature. Specifically, we tested for the effects of birdwatching potential (bird species richness) on NCP that are known to respond to bird diversity: caterpillar predation (Anttonen et al. 2023), acoustic diversity (Müller et al. 2022) and seed predation (Breitbach et al. 2010). We also tested for effects of herbivory on plant pathogen infection, as herbivores can be important vectors of plant pathogens ((Gossner et al. 2021); see *SI Appendix* Table S1).

### Statistical analysis and model selection

Conceptually, our modelling approach predicted the supply of an NCP at a particular plot *i* by accounting for the surrounding landscape (*LUE*_*l,i*_; eq. 1), between-NCP linkages (see ‘NCP data’ above) and other environmental controls (see ‘Landscape explanatory variables’ above). Specifically, we modelled NCP supply *y*_*i*_ using a basic model structure with the set of predictors presented in *SI Appendix* Table S7 for each NCP. At its core, the expected value *E*[*y*_*i*_] is:

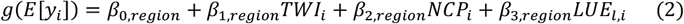

where *g*(·) represents the link function for the corresponding data distribution (*SI Appendix* Table S7), *TWI*_*i*_ is the topographic wetness index of plot *i* that accounts for spatial hydrological processes (Sørensen et al. 2006), *NCP*_*i*_ is another NCP with which *y*_*i*_ may vary, and *LUE*_*l,i*_ is the cumulative spatial land-use effect of land use *l*. We allowed for intercepts and slopes to vary by region (i.e., each *β* has a region subscript). Other components that were added to this basic structure in subsequent models are detailed below.

We applied a model selection approach to determine which model components were important for predicting the supply of each NCP individually (*SI Appendix* Table S2). To do this, we explored various combinations of predictor variables according to our specific hypotheses and assessed model fit using the leave-one-out information criterion (LOOIC) (loo package v2.5.1; (Vehtari et al. 2020)). Firstly, we compared models 1–4 in *SI Appendix* Table S2 to determine whether topographic wetness index (*TWI*_*i*_) and/or relevant other NCP were important for predicting NCP supply and therefore should be included in all subsequent models that explored land-use effects. For each NCP, we then compared models 5–8 with the best-fitting control model in *SI Appendix* Table S2, determining the importance of each land use individually (forest, high-intensity grassland, low-intensity grassland, and cropland). Finally, we ran models that included main effects of land use for pairs of land uses, with all combinations of two-way interactions to test for non-additivity, and compared models with and without interaction terms (models 9– 20 in *SI Appendix* Table S2). We then reran the overall best-fitting models for each NCP with *LUE*_*l,i*_ terms recalculated at all buffer sizes we considered (500, 1000 and 2000 m); comparing parameter estimates confirmed that the choice of buffer size did not qualitatively change our main results (*SI Appendix* Figure S1).

Models were fitted with Bayesian inference using the greta package v0.4.3 (Golding 2019) in R v4.0.4 (R Core Team 2025). We scaled *LUE*_*l,i*_ by dividing it by the maximum effective number of cells (e.g. for a 2000 m buffer, 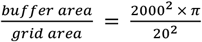). We used Hamiltonian Monte Carlo sampling across four parallel chains, each with 3000 warmup (which were discarded) and 1000 post-warmup samples, which resulted in a total of 4000 posterior samples. All models achieved convergence based on visual assessment of trace plots and the potential scale reduction factor 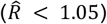 calculated using the coda package v0.19.4 (Plummer et al. 2006). We diagnosed the residuals with Dunn-Smyth randomized quantile residuals (Dunn and Smyth 1996) and quantile-quantile plots. Our data were processed using the sf package v1.0.5 (Pebesma 2018) and visualized using the ggplot2 v3.4.1 (Wickham 2009) and bayesplot v1.10.0 packages (Gabry et al. 2019).

### Calculation of netNCP

We hypothesized that the supply of multiple NCP will vary spatially and generate trade-offs due to varying responses of NCP to different land uses in the surrounding landscape. We visualized these trade-offs as the weighted sum of beneficial and detrimental NCP (netNCP; (Neyret et al. 2024)). In this calculation, NCP were weighted such that beneficial and detrimental NCP were equally weighted by dividing by the total number of beneficial (five) or detrimental (three) NCP respectively.

## Acknowledgments

S.A.O’B. was supported by a Food Transitions 2050 Doctoral Scholarship (University of Canterbury), a William Georgetti Scholarship ((New Zealand Vice Chancellor’s Committee) and the Edward and Isabel Kidson Scholarship (New Zealand Vice Chancellor’s Committee). Funding for this work was provided in part by the Ministry of Business, Innovation & Employment (program number C09×2209), and J.M.T. and H.R.L are funded by the Bioprotection Aotearoa Centre of Research Excellence. H.R.L. is supported by the Marsden Fund managed by the Royal Society Te Apārangi (grant MFP-UOC2102). S.L. acknowledges support by Biodiversa+ project RECONNECT through Agence Nationale pour la Recherche (ANR-22-EBIP-0009-06). The authors wish to acknowledge the use of New Zealand eScience Infrastructure (NeSI) high performance computing facilities and consulting support as part of this research. New Zealand’s national facilities are provided by NeSI and funded jointly by NeSI’s collaborator institutions and through the Ministry of Business, Innovation & Employment’s Research Infrastructure program, http://www.nesi.org.nz.

The work has been partly funded by the DFG Priority Program 1374 “Biodiversity Exploratories”. Field work permits were issued by the responsible state environmental offices of Baden-Württemberg, Thüringen, and Brandenburg. We thank K. Wells, K. Reichel-Jung, S. Gockel, K. Wiesner, K. Lorenzen, A. Hemp, M. Gorke for their work in maintaining the plot and project infrastructure; S. Pfeiffer, M. Gleisberg, C. Fischer, J. Mangels, and V. Grießmeier for giving support through the central office, J. Nieschulze, M. Owonibi, and A. Ostrowski for database management, and M. Fischer, E. Linsenmair, D. Hessenmöller, D. Prati, I. Schöning, E.-D. Schulze, W. W. Weisser, and E. Kalko for their role in setting up the Biodiversity Exploratories project. We thank the administration of the Hainich National Park, the UNESCO Biosphere Reserves Swabian Alb and Schorfheide-Chorin as well as all land owners for the excellent collaboration. Thanks to Eric Allan for data input.

## Data Availability

Most datasets and associated metadata are publicly available on the Biodiversity Exploratories Information System at https://doi.org/10.17616/R32P9Q (IDs: 21447, 27568, 27569, 27570, 25806, 24966). The remaining datasets (IDs: 27087, 27727, 31018) are subject to an embargo period of three years, and will be made available at the same data repository at the conclusion of the respective embargo period. Correspondence for specific datasets should be directed to the respective data owners listed in *SI Appendix* Table S6. Code used to prepare data and conduct analyses will be made available in the same repository.

## References

Adem Esmail, B., and D. Geneletti. 2018. Multi-criteria decision analysis for nature conservation: A review of 20 years of applications. Methods in Ecology and Evolution 9:42–53.

Anttonen, P., M. Perles-Garcia, M. Kunz, G. von Oheimb, Y. Li, H. Bruelheide, K.-P. Ma, C.-D. Zhu, and A. Schuldt. 2023. Predation pressure by arthropods, birds, and rodents is interactively shaped by tree species richness, vegetation structure, and season. Frontiers in Ecology and Evolution 11:1199670.

Blitzer, E. J., C. F. Dormann, A. Holzschuh, A.-M. Klein, T. A. Rand, and T. Tscharntke. 2012. Spillover of functionally important organisms between managed and natural habitats. Agriculture, Ecosystems & Environment 146:34–43.

Boesing, A. L., V. H. Klaus, M. Neyret, G. Le Provost, S. Peter, M. Fischer, and P. Manning. 2024. Identifying the optimal landscape configuration for landscape multifunctionality. Ecosystem services 67:101630.

Breitbach, N., I. Laube, I. Steffan-Dewenter, and K. Böhning-Gaese. 2010. Bird diversity and seed dispersal along a human land-use gradient: high seed removal in structurally simple farmland. Oecologia 162:965–976.

Cholewa, M., and T. Wesołowski. 2011. Nestling food of European hole-nesting passerines: do we know enough to test the adaptive hypotheses on breeding seasons? Acta Ornithologica 46:105–116.

Clark, J. S., M. Silman, R. Kern, E. Macklin, and J. HilleRisLambers. 1999. Seed dispersal near and far: patterns across temperate and tropical forests. Ecology 80:1475–1494.

Convention on Biological Diversity. 2022. Kunming-Montreal Global Biodiversity Framework. https://www.cbd.int/doc/decisions/cop-15/cop-15-dec-04-en.pdf.

Dade, M. C., M. G. E. Mitchell, C. A. McAlpine, and J. R. Rhodes. 2019. Assessing ecosystem service trade-offs and synergies: The need for a more mechanistic approach. Ambio 48:1116–1128.

Díaz, S., U. Pascual, M. Stenseke, B. Martín-López, R. T. Watson, Z. Molnár, R. Hill, K. M. A. Chan, I. A. Baste, and K. A. Brauman. 2018. Assessing nature’s contributions to people. Science 359:270– 272.

Dunn, P. K., and G. K. Smyth. 1996. Randomized quantile residuals. Journal of Computational and graphical statistics 5:236–244.

Fischer, C., C. Gayer, K. Kurucz, F. Riesch, T. Tscharntke, and P. Batáry. 2018. Ecosystem services and disservices provided by small rodents in arable fields: Effects of local and landscape management. Journal of applied ecology 55:548–558.

Fischer, M., O. Bossdorf, S. Gockel, F. Hänsel, A. Hemp, D. Hessenmöller, G. Korte, J. Nieschulze, S. Pfeiffer, and D. Prati. 2010. Implementing large-scale and long-term functional biodiversity research: The Biodiversity Exploratories. Basic and Applied Ecology 11:473–485.

Gabry, J., D. Simpson, A. Vehtari, M. Betancourt, and A. Gelman. 2019. Visualization in Bayesian workflow. Journal of the Royal Statistical Society Series A: Statistics in Society 182:389–402.

Garibaldi, L. A., I. Steffan-Dewenter, C. Kremen, J. M. Morales, R. Bommarco, S. A. Cunningham, L. G. Carvalheiro, N. P. Chacoff, J. H. Dudenhöffer, and S. S. Greenleaf. 2011. Stability of pollination services decreases with isolation from natural areas despite honey bee visits. Ecology Letters 14:1062–1072.

Golding, N. 2019. greta: simple and scalable statistical modelling in R. Journal of Open Source Software 4:1601.

Gomes, E., M. Inácio, K. Bogdzevič, M. Kalinauskas, D. Karnauskaitė, and P. Pereira. 2021. Future landuse changes and its impacts on terrestrial ecosystem services: A review. Science of The Total Environment 781:146716.

Gossner, M. M., L. Beenken, K. Arend, D. Begerow, and D. Peršoh. 2021. Insect herbivory facilitates the establishment of an invasive plant pathogen. ISME communications 1:6.

Han, Y., Y. Liu, X. Wu, and Q. Zhang. 2024. Assessment and forecast of the water-related nature’s contributions to people on the Loess Plateau from a spatial flow perspective. Landscape Ecology 39:159.

Hohlenwerger, C., R. Spake, L. R. Tambosi, N. Aristizábal, A. González-Chaves, F. Librán-Embid, F. Saturni, F. Eigenbrod, and J.-P. Metzger. 2024. Coffee pollination and pest control are affected by edge diversity at local scales but multiscalar approaches and disservices can not be ignored. Landscape Ecology 39:75.

Jenkins, D. G., C. R. Brescacin, C. V. Duxbury, J. A. Elliott, J. A. Evans, K. R. Grablow, M. Hillegass, B. N. Lyon, G. A. Metzger, and M. L. Olandese. 2007. Does size matter for dispersal distance? Global Ecology and Biogeography 16:415–425.

Kass, J. M., K. Fukaya, W. Thuiller, and A. S. Mori. 2024. Biodiversity modeling advances will improve predictions of nature’s contributions to people. Trends in Ecology & Evolution 39:338–348.

Lander, T. A., D. P. Bebber, C. T. Choy, S. A. Harris, and D. H. Boshier. 2011. The Circe principle explains how resource-rich land can waylay pollinators in fragmented landscapes. Current Biology 21:1302–1307.

Lange, M., H. Feilhauer, I. Kühn, and D. Doktor. 2022. Mapping land-use intensity of grasslands in Germany with machine learning and Sentinel-2 time series. Remote Sensing of Environment 277:112888.

Le Provost, G., N. V. Schenk, C. Penone, J. Thiele, C. Westphal, E. Allan, M. Ayasse, N. Blüthgen, R. S. Boeddinghaus, A. L. Boesing, and et al. 2023. The supply of multiple ecosystem services requires biodiversity across spatial scales. Nature ecology & evolution 7:236–249.

Magdon, P. 2023. Land cover (LBM-DE) of all Biodiversity Exploratories regions. Biodiversity Exploratories Information System, https://www.bexis.uni-jena.de/ddm/data/Showdata/27727.

Manning, P. 2023. Aggregated environmental and land-use covariates of the 150 grassland EPs used in “Contrasting responses of above- and belowground diversity to multiple components of land-use intensity”. Biodiversity Exploratories Information System, https://www.bexis.uni-jena.de/ddm/data/Showdata/31018?version=5.

Metzger, J. P., J. Villarreal-Rosas, A.F. Suárez-Castro, S. López-Cubillos, A. González-Chaves, R. K. Runting, C. Hohlenwerger, and J. R. Rhodes. 2021. Considering landscape-level processes in ecosystem service assessments. Science of The Total Environment 796:149028.

Mohamed, A., F. DeClerck, P. H. Verburg, D. Obura, J. F. Abrams, N. Zafra-Calvo, J. Rocha, N. Estrada-Carmona, A. Fremier, S. K. Jones, I. C. Meier, and B. Stewart-Koster. 2024. Securing Nature’s Contributions to People requires at least 20%–25% (semi-)natural habitat in human-modified landscapes. One Earth 7:59–71.

Müller, S., M. M. Gossner, C. Penone, K. Jung, S. C. Renner, A. Farina, L. Anhäuser, M. Ayasse, S. Boch, and F. Haensel. 2022. Land-use intensity and landscape structure drive the acoustic composition of grasslands. Agriculture, Ecosystems & Environment 328:107845.

Newbold, T., J. P. Scharlemann, S. H. Butchart, Ç. H. Sekercioglu, R. Alkemade, H. Booth, and D. W. Purves. 2013. Ecological traits affect the response of tropical forest bird species to land-use intensity. Proceedings of the Royal Society B: Biological Sciences 280:20122131.

Neyret, M., A. L. Boesing, G. Le Provost, S. Peter, N. R. Kinabo, D. G. Mauki, K. D. Koggani, A. Thiel, B. Martín-López, and P. Manning. 2024. Expanding ecosystem multifunctionality measures to operationalize the IPBES framework. Preprint:https://hal.science/hal-04390432v04390431.

Neyret, M., M. Fischer, E. Allan, N. Hölzel, V. H. Klaus, T. Kleinebecker, J. Krauss, G. Le Provost, S. Peter, N. Schenk, N. K. Simons, F. van der Plas, J. Binkenstein, C. Börschig, K. Jung, D. Prati, D. Schäfer, M. Schäfer, I. Schöning, M. Schrumpf, M. Tschapka, C. Westphal, and P. Manning. 2021. Assessing the impact of grassland management on landscape multifunctionality. Ecosystem services 52:101366.

O’Brien, S. A., D. P. Anderson, S. Lavorel, H. R. Lai, N. de Burgh, and J. M. Tylianakis. 2024. Landscape patterns drive provision of nature’s contributions to people by mobile species. Journal of applied ecology 61:2666–2678.

Pebesma, E. J. 2018. Simple features for R: standardized support for spatial vector data. R J. 10:439.

Peña, R., M. Schleuning, M. Miñarro, and D. García. 2023. Variable relationships between trait diversity and avian ecological functions in agroecosystems. Functional ecology 37:87–98.

Plummer, M., N. Best, K. Cowles, and K. Vines. 2006. CODA: convergence diagnosis and output analysis for MCMC. R news 6:7–11.

R Core Team, R. 2025. R: A language and environment for statistical computing.

Rey, P.-L., C. Martin, and A. Guisan. 2024. Conservation importance of non-threatened species through their direct linkages with nature’s contributions to people. Biological Conservation 297:110733.

Ricketts, T. H., J. Regetz, I. Steffan-Dewenter, S. A. Cunningham, C. Kremen, A. Bogdanski, B. Gemmill-Herren, S. S. Greenleaf, A. M. Klein, M. M. Mayfield, L. A. Morandin, A. Ochieng’, and B. F. Viana. 2008. Landscape effects on crop pollination services: are there general patterns? Ecology Letters 11:499–515.

Seidel, D. P., E. Dougherty, C. Carlson, and W. M. Getz. 2018. Ecological metrics and methods for GPS movement data. International Journal of Geographical Information Science 32:2272–2293.

Silveira, N. S. D., B. B. S. Niebuhr, R. de Lara Muylaert, M. C. Ribeiro, and M. A. Pizo. 2016. Effects of land cover on the movement of frugivorous birds in a heterogeneous landscape. PloS one 11:1– 19.

Simons, N. K., M. R. Felipe-Lucia, P. Schall, C. Ammer, J. Bauhus, N. Blüthgen, S. Boch, F. Buscot, M. Fischer, K. Goldmann, M. M. Gossner, F. Hänsel, K. Jung, P. Manning, T. Nauss, Y. Oelmann, R. Pena, A. Polle, S. C. Renner, M. Schloter, I. Schöning, E.-D. Schulze, E. F. Solly, E. Sorkau, B. Stempfhuber, T. Wubet, J. Müller, S. Seibold, and W. W. Weisser. 2021. National Forest Inventories capture the multifunctionality of managed forests in Germany. Forest Ecosystems 8:5.

Sørensen, R., U. Zinko, and J. Seibert. 2006. On the calculation of the topographic wetness index: evaluation of different methods based on field observations. Hydrology and Earth System Sciences 10:101–112.

Spake, R., R. Lasseur, E. Crouzat, J. M. Bullock, S. Lavorel, K. E. Parks, M. Schaafsma, E. M. Bennett, J. Maes, and M. Mulligan. 2017. Unpacking ecosystem service bundles: Towards predictive mapping of synergies and trade-offs between ecosystem services. Global Environmental Change 47:37–50.

Stein, A., K. Gerstner, and H. Kreft. 2014. Environmental heterogeneity as a universal driver of species richness across taxa, biomes and spatial scales. Ecology Letters 17:866–880.

Vanneste, T., L. Depauw, E. De Lombaerde, C. Meeussen, S. Govaert, K. De Pauw, P. Sanczuk, K. Bollmann, J. Brunet, and K. Calders. 2024. Trade-offs in biodiversity and ecosystem services between edges and interiors in European forests. Nature ecology & evolution 8:880–887.

Vehtari, A., J. Gabry, M. Magnusson, Y. Yao, P.-C. Bürkner, T. Paananen, and A. Gelman. 2020. loo: Efficient leave-one-out cross-validation and WAIC for Bayesian models. R package version 2:12.

Wehner, K., L. Schäfer, N. Blüthgen, and K. Mody. 2020. Seed type, habitat and time of day influence post-dispersal seed removal in temperate ecosystems. PeerJ 8:e8769.

Wickham, H. 2009. ggplot2: elegant graphics for data analysis. Springer, New York.

