## Supporting Information for "Mobile species’ responses to surrounding land use generate trade-offs among nature’s contributions to people"

**This PDF file includes:**

Supporting text
Fig. S1
Tables S1 to S7
SI References

### Supporting Information Text

#### Sensitivity analysis

The majority of results were robust to the scale at which land use effects ($LUE_{l,i}$) were calculated (*SI Appendix* Fig. S1). Specifically, 90.4 % (85) of the coefficients estimated did not significantly change when land-use effects were recalculated at 500, 1000 and 2000 buffers (i.e. the result presented in the main text fell within the 90 % credible intervals of analyses using differently-scaled buffers). The remaining 9 coefficient estimates that did show some scale-dependent responses were limited to two NCP: pollination and natural enemy abundance. Four of these coefficient estimates changed significantly in magnitude across scales. One coefficient estimate showed a significant change in the sign, but the 90 % credible intervals overlapped with zero. Finally, for 4 out of 94 parameters estimated (4.3 %), all from the models of pollination, there was a significant change in both the sign and magnitude of the coefficient estimates as the spatial scale of measurement changed. This aligns with previous research that found scale-dependent pollinator responses to landscape composition and configuration; key processes acting on pollinator species operate at different spatial scales, such as those related to availability of floral resources and interaction partners (1, 2).

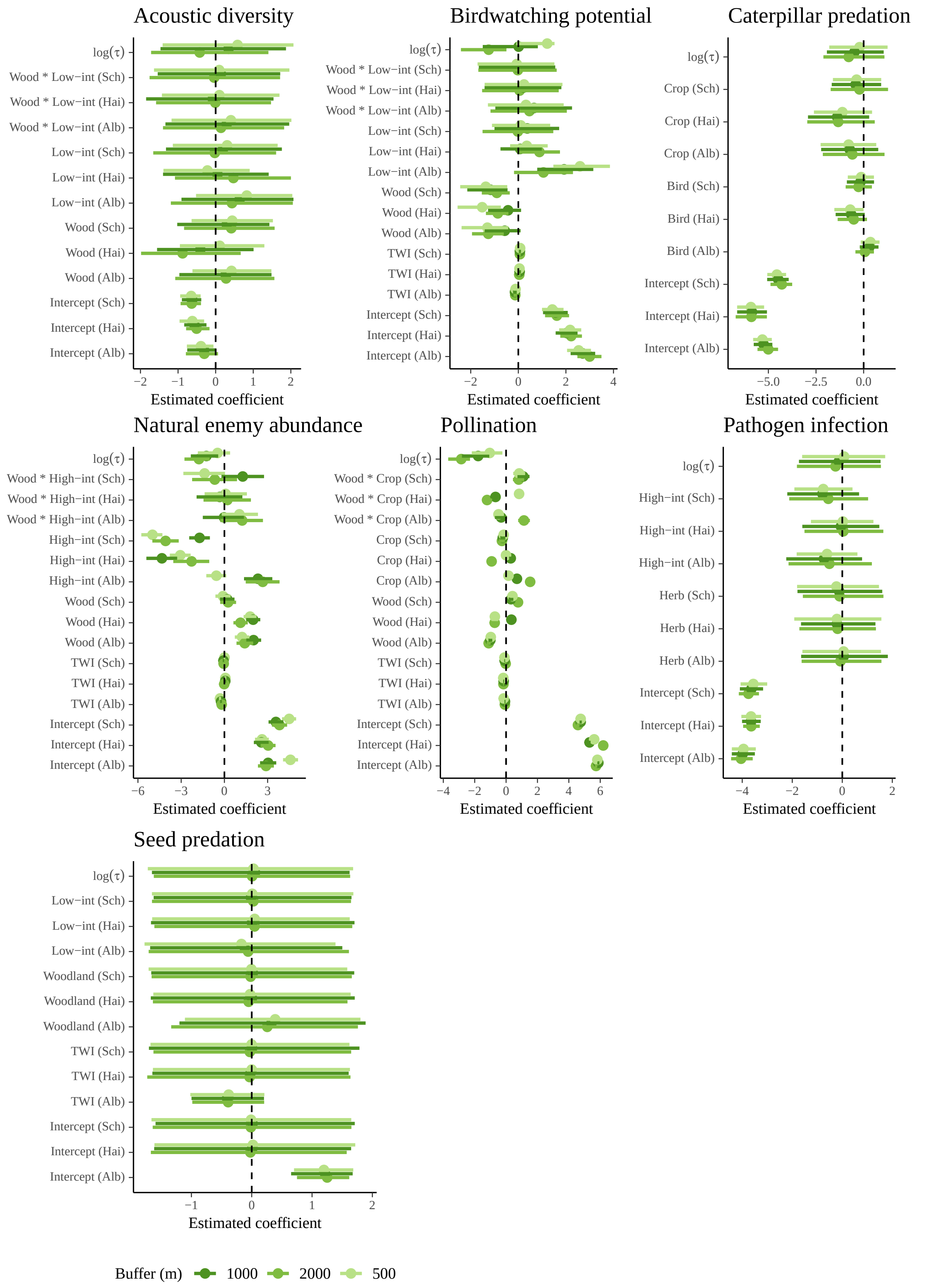


**Fig. S1.** Parameter estimates from our main results for each NCP (the best-fitting models in *SI Appendix* Table S2) presented with those same models rerun with $\mathrm{LUE}_{l,i}$ calculated using a range of buffer sizes (500, 1000 and 2000 m), including those used in main analyses (see *SI Appendix* Table S5). $\log(\tau)$ is the inverse of $\log(\gamma)$, the distance-decay parameter used in equations 1 and 2 in 'Materials and Methods.' Parameter abbreviations are as follows: low-int is low-intensity grassland; high-int is high-intensity grassland; wood is woodland; crop is cropland; bird is birdwatching potential and herb is herbivory. Alb, Hai and Sch correspond to the region fixed effects for Schwäbische Alb, Hainich-Dün and Schorfheide-Chorin respectively. Points and error bars are posterior median and 90% credible intervals, respectively. The NCP herbivory is omitted from this figure because the best-fit model did not retain any $\mathrm{LUE}_{l,i}$ for which we could test the sensitivity of results to buffer size.

**Table S1. Hypothesized responses of NCP to land use, and relationships with potential NCP covariates.**

| NCP | Beneficial or detrimental | Hypothesized response to land use surrounding grassland plots | NCP covariates | Mechanism | Reference |
|---|---|---|---|---|---|
| Acoustic diversity | Beneficial | Higher in grassland than forest (3) | Birdwatching potential | Higher bird species richness contributes to greater diversity of acoustic bands | (4) |
| Birdwatching potential | Beneficial | Higher in grassland than forest (3) | – | – | – |
| Caterpillar predation | Beneficial | Higher in land use with highest bird species richness (5) | Birdwatching potential | Caterpillar predation by birds is higher with increased bird species richness (which in Europe is higher in grasslands) | (3, 5) |
| Natural enemy abundance | Beneficial | Higher in low-intensity than high-intensity grassland (6) | – | – | – |
| Pollination | Beneficial | Higher in forest and low-intensity grassland (7) | – | – | – |
| Herbivory | Detrimental | Higher in cropland (8) | – | – | – |
| Pathogen infection | Detrimental | Lowest in land use with highest plant richness via mediation of infection pressure (9) | Herbivory | Herbivory facilitates pathogen infection by creating entry points and favorable conditions | (10) |
| Seed predation | Detrimental | Higher in grassland than forest (11) | Birdwatching potential | Seed predation is higher with increased bird species richness | (12) |

**Table S2. Full list of models tested, with their model fit relative to the best-fitting model (difference in Leave-one-out information criterion; △LOOIC), and parameters included in predicting NCP provision. For all models, the proportion of parameters for which $\hat{R} < 1.05$ was $> 0.95$ (a measure of convergence). NCP abbreviations are as follows: ADI is acoustic diversity; BIRD is birdwatching potential; CATER is caterpillar predation; NATEN is natural enemy abundance; POLL is pollination; HERB is herbivory; PATHO is pathogen infection; SEED is seed predation. The △LOOIC value of the best-fitting controls model is italicized, corresponding to the control variables that were included in all subsequent models. The control variables giving the best model fit varied by NCP and are as follows: Intercept only: ADI, HERB; Intercept + TWI: BIRD, NATEN, POLL, SEED; Intercept + BIRD: CATER; Intercept + HERB: PATHO. The △LOOIC value of the best-fitting overall model (i.e. 0) is in bold. LUE (land-use effects) suffixes correspond to the following land uses: low-int is low-intensity grassland; high-int is high-intensity grassland.**

| Model | Parameters predicting NCP provision | Beneficial | | | | | Detrimental | | |
|---|---|---|---|---|---|---|---|---|---|
| | | ADI | BIRD | CATER | NATEN | POLL | HERB | PATHO | SEED |
| | NCP covariate tested | BIRD | – | BIRD | – | – | – | HERB | BIRD |
| 1 | $\beta_0$ | *2.1* | 51.9 | 135.0 | 397.0 | 1156.7 | ***0*** | 0.2 | 8622.0 |
| 2 | $\beta_0 + \beta_1 \text{TWI}_i$ | 5.3 | *34.9* | 148.7 | *355.2* | *900.4* | 2.8 | 7.9 | *8216.8* |
| 3 | $\beta_0 + \beta_1 \text{NCP}_i$ | 5.5 | – | *132.4* | – | – | – | *0.04* | 8486.3 |
| 4 | $\beta_0 + \beta_1 \text{TWI}_i + \beta_2 \text{NCP}_i$ | 9.3 | – | 149.2 | – | – | – | 7.6 | 8307.0 |
| 5 | $\beta_0 + \beta_1 \text{LUE}_{\text{forest},i} + \text{Controls}$ | 0.5 | 2.5 | 59.6 | 303.9 | 454.2 | 2.1 | 1.5 | 6.2 |
| 6 | $\beta_0 + \beta_1 \text{LUE}_{\text{low-int},i} + \text{Controls}$ | 1.7 | 29.8 | 21.5 | 337.6 | 934.8 | 1.1 | 1.0 | 12.1 |
| 7 | $\beta_0 + \beta_1 \text{LUE}_{\text{high-int},i} + \text{Controls}$ | 1.8 | 26.0 | 9.1 | 60.2 | 945.0 | 1.5 | **0** | 17.4 |
| 8 | $\beta_0 + \beta_1 \text{LUE}_{\text{cropland},i} + \text{Controls}$ | 1.6 | 23.2 | **0** | 293.5 | 306.5 | 1.7 | 0.8 | 8.8 |
| 9 | $\beta_0 + \beta_1 \text{LUE}_{\text{forest},i} + \beta_2 \text{LUE}_{\text{low-int},i} + \text{Controls}$ | 0.7 | 1.1 | 58.8 | 234.8 | 271.2 | 2.1 | 1.5 | **0** |
| 10 | $\beta_0 + \beta_1 \text{LUE}_{\text{forest},i} + \beta_2 \text{LUE}_{\text{high-int},i} + \text{Controls}$ | 0.8 | 0.6 | 20.0 | 9.9 | 471.9 | 1.3 | 0.3 | 12.0 |
| 11 | $\beta_0 + \beta_1 \text{LUE}_{\text{forest},i} + \beta_2 \text{LUE}_{\text{cropland},i} + \text{Controls}$ | 0.7 | 0.5 | 3.8 | 218.1 | 86.4 | 2.4 | 0.3 | 7.6 |
| 12 | $\beta_0 + \beta_1 \text{LUE}_{\text{low-int},i} + \beta_2 \text{LUE}_{\text{high-int},i} + \text{Controls}$ | 1.4 | 0.6 | 62.9 | 66.3 | 959.3 | 1.1 | 1.9 | 10.8 |
| 13 | $\beta_0 + \beta_1 \text{LUE}_{\text{low-int},i} + \beta_2 \text{LUE}_{\text{cropland},i} + \text{Controls}$ | 1.4 | 1.2 | 19.8 | 308.7 | 360.6 | 1.4 | 0.5 | 15.3 |
| 14 | $\beta_0 + \beta_1 \text{LUE}_{\text{high-int},i} + \beta_2 \text{LUE}_{\text{cropland},i} + \text{Controls}$ | 1.3 | 0.9 | 8.1 | 22.7 | 375.4 | 1.4 | 0.5 | 4.6 |
| 15 | $\beta_0 + \beta_1 \text{LUE}_{\text{forest},i} + \beta_2 \text{LUE}_{\text{low-int},i} + \beta_3 \text{LUE}_{\text{forest},i} \times \text{LUE}_{\text{low-int},i} + \text{Controls}$ | **0** | **0** | 59.1 | 232.1 | 266.5 | 1.7 | 1.3 | 7.0 |
| 16 | $\beta_0 + \beta_1 \text{LUE}_{\text{forest},i} + \beta_2 \text{LUE}_{\text{high-int},i} + \beta_3 \text{LUE}_{\text{forest},i} \times \text{LUE}_{\text{high-int},i} + \text{Controls}$ | 0.7 | 0.6 | 20.7 | **0** | 501.2 | 2.0 | 0.6 | 6.5 |
| 17 | $\beta_0 + \beta_1 \text{LUE}_{\text{forest},i} + \beta_2 \text{LUE}_{\text{cropland},i} + \beta_3 \text{LUE}_{\text{forest},i} \times \text{LUE}_{\text{cropland},i} + \text{Controls}$ | 0.8 | 6.2 | 3.5 | 212.6 | **0** | 2.5 | 0.5 | 17.2 |
| 18 | $\beta_0 + \beta_1 \text{LUE}_{\text{low-int},i} + \beta_2 \text{LUE}_{\text{high-int},i} + \beta_3 \text{LUE}_{\text{low-int},i} \times \text{LUE}_{\text{high-int},i} + \text{Controls}$ | 1.3 | 10.6 | 62.0 | 61.3 | 935.5 | 1.6 | 0.05 | 7.6 |
| 19 | $\beta_0 + \beta_1 \text{LUE}_{\text{low-int},i} + \beta_2 \text{LUE}_{\text{cropland},i} + \beta_3 \text{LUE}_{\text{low-int},i} \times \text{LUE}_{\text{cropland},i} + \text{Controls}$ | 1.1 | 4.9 | 20.2 | 275.9 | 360.9 | 2.1 | 1.0 | 17.0 |
| 20 | $\beta_0 + \beta_1 \text{LUE}_{\text{high-int},i} + \beta_2 \text{LUE}_{\text{cropland},i} + \beta_3 \text{LUE}_{\text{high-int},i} \times \text{LUE}_{\text{cropland},i} + \text{Controls}$ | 1.2 | 16.5 | 7.8 | 13.5 | 372.7 | 1.6 | 0.4 | 8.7 |

**Table S3. Posterior means and $90\,\%$ credible intervals of model coefficients for the best-fitting model in *SI Appendix* Table S2 for each NCP, by region. LUE suffixes low-int and high-int correspond to low- and high-intensity grassland respectively. $\text{Bird}_i$ corresponds to birdwatching potential and $\text{Herb}_i$ to herbivory. Regions are Schwäbische Alb (Alb); Hainich-Dün (Hai); and Schorfheide-Chorin (Sch).**

| NCP | Coefficient | Variable | Region | Mean | 5 % CI | 95 % CI |
|---|---|---|---|---|---|---|
| Beneficial | | | | | | |
| Acoustic diversity | $\beta_0$ | Intercept | Alb | -0.327 | -0.792 | 0.063 |
| | $\beta_0$ | | Hai | -0.494 | -0.786 | -0.163 |
| | $\beta_0$ | | Sch | -0.647 | -0.933 | -0.390 |
| | $\beta_1$ | $\text{LUE}_{\text{forest},i}$ | Alb | 0.261 | -1.075 | 1.563 |
| | $\beta_1$ | | Hai | -0.802 | -1.983 | 0.666 |
| | $\beta_1$ | | Sch | 0.394 | -0.840 | 1.568 |
| | $\beta_2$ | $\text{LUE}_{\text{low-int},i}$ | Alb | 0.434 | -1.192 | 2.053 |
| | $\beta_2$ | | Hai | 0.475 | -1.082 | 2.003 |
| | $\beta_2$ | | Sch | -0.018 | -1.660 | 1.608 |
| | $\beta_3$ | $\text{LUE}_{\text{forest},i} \times \text{LUE}_{\text{low-int},i}$ | Alb | 0.164 | -1.401 | 1.822 |
| | $\beta_3$ | | Hai | -0.034 | -1.585 | 1.473 |
| | $\beta_3$ | | Sch | -0.040 | -1.758 | 1.714 |
| | $\gamma$ | | | 2.083 | 0.246 | 5.568 |
| Birdwatching potential | $\beta_0$ | Intercept | Alb | 2.996 | 2.483 | 3.495 |
| | $\beta_0$ | | Hai | 2.222 | 1.764 | 2.674 |
| | $\beta_0$ | | Sch | 1.622 | 1.123 | 2.133 |
| | $\beta_1$ | $\text{TWI}_i$ | Alb | -0.128 | -0.200 | -0.058 |
| | $\beta_1$ | | Hai | 0.040 | -0.014 | 0.093 |
| | $\beta_1$ | | Sch | 0.065 | 0.022 | 0.107 |
| | $\beta_2$ | $\text{LUE}_{\text{forest},i}$ | Alb | -1.267 | -1.946 | -0.609 |
| | $\beta_2$ | | Hai | -0.871 | -1.362 | -0.415 |
| | $\beta_2$ | | Sch | -0.914 | -1.534 | -0.358 |
| | $\beta_3$ | $\text{LUE}_{\text{low-int},i}$ | Alb | 1.060 | -0.180 | 2.297 |
| | $\beta_3$ | | Hai | 0.878 | -0.057 | 1.751 |
| | $\beta_3$ | | Sch | -0.019 | -1.504 | 1.470 |
| | $\beta_4$ | $\text{LUE}_{\text{forest},i} \times \text{LUE}_{\text{low-int},i}$ | Alb | 0.446 | -1.169 | 2.038 |
| | $\beta_4$ | | Hai | 0.060 | -1.522 | 1.698 |
| | $\beta_4$ | | Sch | -0.039 | -1.681 | 1.613 |
| | $\gamma$ | | | 4.556 | 1.647 | 11.154 |
| Caterpillar predation | $\beta_0$ | Intercept | Alb | -5.315 | -5.800 | -4.819 |
| | $\beta_0$ | | Hai | -5.927 | -6.641 | -5.224 |
| | $\beta_0$ | | Sch | -4.558 | -5.058 | -4.070 |
| | $\beta_1$ | $\text{Bird}_i$ | Alb | 0.347 | -0.153 | 0.833 |
| | $\beta_1$ | | Hai | -0.729 | -1.537 | -0.006 |
| | $\beta_1$ | | Sch | -0.140 | -0.833 | 0.541 |
| | $\beta_2$ | $\text{LUE}_{\text{cropland},i}$ | Alb | -0.784 | -2.258 | 0.662 |
| | $\beta_2$ | | Hai | -1.120 | -2.610 | 0.442 |
| | $\beta_2$ | | Sch | -0.370 | -1.613 | 0.923 |
| | $\gamma$ | | | 1.988 | 0.285 | 6.084 |
| Natural enemy abundance | $\beta_0$ | Intercept | Alb | 4.584 | 4.069 | 5.109 |
| | $\beta_0$ | | Hai | 2.608 | 2.123 | 3.081 |
| | $\beta_0$ | | Sch | 4.493 | 3.998 | 4.976 |
| | $\beta_1$ | $\text{TWI}_i$ | Alb | -0.304 | -0.379 | -0.231 |
| | $\beta_1$ | | Hai | 0.068 | 0.004 | 0.132 |
| | $\beta_1$ | | Sch | 0.007 | -0.046 | 0.057 |
| | $\beta_2$ | $\text{LUE}_{\text{forest},i}$ | Alb | 1.229 | 0.723 | 1.739 |
| | $\beta_2$ | | Hai | 1.783 | 1.331 | 2.227 |
| | $\beta_2$ | | Sch | -0.096 | -0.624 | 0.428 |
| | $\beta_3$ | $\text{LUE}_{\text{high-int},i}$ | Alb | -0.565 | -1.256 | 0.129 |
| | $\beta_3$ | | Hai | -3.060 | -3.787 | -2.343 |
| | $\beta_3$ | | Sch | -5.012 | -5.765 | -4.302 |
| | $\beta_4$ | $\text{LUE}_{\text{forest},i} \times \text{LUE}_{\text{high-int},i}$ | Alb | 1.044 | -0.185 | 2.329 |
| | $\beta_4$ | | Hai | 0.075 | -1.374 | 1.559 |
| | $\beta_4$ | | Sch | -1.385 | -2.848 | 0.040 |
| | $\gamma$ | | | 2.368 | 0.674 | 6.323 |
| Pollination | $\beta_0$ | Intercept | Alb | 5.736 | 5.661 | 5.811 |
| | $\beta_0$ | | Hai | 6.193 | 6.088 | 6.300 |
| | $\beta_0$ | | Sch | 4.580 | 4.487 | 4.677 |
| | $\beta_1$ | $\text{TWI}_i$ | Alb | -0.075 | -0.107 | -0.043 |
| | $\beta_1$ | | Hai | -0.168 | -0.197 | -0.140 |
| | $\beta_1$ | | Sch | -0.027 | -0.057 | 0.002 |
| | $\beta_2$ | $\text{LUE}_{\text{forest},i}$ | Alb | -1.109 | -1.234 | -0.982 |
| | $\beta_2$ | | Hai | -0.726 | -0.888 | -0.570 |
| | $\beta_2$ | | Sch | 0.760 | 0.587 | 0.930 |
| | $\beta_3$ | $\text{LUE}_{\text{cropland},i}$ | Alb | 1.532 | 1.295 | 1.768 |
| | $\beta_3$ | | Hai | -0.921 | -1.073 | -0.778 |
| | $\beta_3$ | | Sch | -0.266 | -0.462 | -0.073 |
| | $\beta_4$ | $\text{LUE}_{\text{forest},i} \times \text{LUE}_{\text{cropland},i}$ | Alb | 1.141 | 0.768 | 1.516 |
| | $\beta_4$ | | Hai | -1.214 | -1.536 | -0.891 |
| | $\beta_4$ | | Sch | 0.801 | 0.437 | 1.154 |
| | $\gamma$ | | | 20.222 | 9.969 | 39.997 |

Continued on next page

**Table S3 – continued from previous page**

| NCP | Coefficient | Variable | Region | Mean | 5 % CI | 95 % CI |
|---|---|---|---|---|---|---|
| Detrimental | | | | | | |
| Herbivory | $\beta_0$ | Intercept | Alb | -3.920 | -4.140 | -3.702 |
| | $\beta_0$ | | Hai | -3.663 | -3.867 | -3.470 |
| | $\beta_0$ | | Sch | -3.492 | -3.680 | -3.300 |
| Pathogen infection | $\beta_0$ | Intercept | Alb | -4.033 | -4.445 | -3.580 |
| | $\beta_0$ | | Hai | -3.636 | -3.964 | -3.299 |
| | $\beta_0$ | | Sch | -3.741 | -4.136 | -3.331 |
| | $\beta_1$ | $\text{Herb}_i$ | Alb | -0.048 | -1.626 | 1.566 |
| | $\beta_1$ | | Hai | -0.190 | -1.716 | 1.345 |
| | $\beta_1$ | | Sch | -0.019 | -1.577 | 1.646 |
| | $\beta_2$ | $\text{LUE}_{\text{high-int},i}$ | Alb | -0.504 | -2.148 | 1.183 |
| | $\beta_2$ | | Hai | 0.047 | -1.511 | 1.641 |
| | $\beta_2$ | | Sch | -0.545 | -2.122 | 1.033 |
| | $\gamma$ | | | 1.984 | 0.213 | 6.135 |
| Seed predation | $\beta_0$ | Intercept | Alb | 1.229 | 0.750 | 1.617 |
| | $\beta_0$ | | Hai | -0.025 | -1.672 | 1.575 |
| | $\beta_0$ | | Sch | -0.002 | -1.641 | 1.653 |
| | $\beta_1$ | $\text{TWI}_i$ | Alb | -0.393 | -0.985 | 0.206 |
| | $\beta_1$ | | Hai | -0.017 | -1.732 | 1.637 |
| | $\beta_1$ | | Sch | -0.013 | -1.629 | 1.649 |
| | $\beta_2$ | $\text{LUE}_{\text{forest},i}$ | Alb | 0.223 | -1.335 | 1.761 |
| | $\beta_2$ | | Hai | -0.037 | -1.638 | 1.588 |
| | $\beta_2$ | | Sch | -0.013 | -1.660 | 1.659 |
| | $\beta_3$ | $\text{LUE}_{\text{low-int},i}$ | Alb | -0.075 | -1.708 | 1.611 |
| | $\beta_3$ | | Hai | 0.020 | -1.614 | 1.667 |
| | $\beta_3$ | | Sch | 0.022 | -1.654 | 1.646 |
| | $\gamma$ | | | 1.644 | 0.196 | 5.070 |

**Table S4. The relative percentages of each land use in our study regions, with increasingly broad categorization. Region abbreviations are as follows: Alb is Schwäbische Alb; Hai is Hainich-Dün; Sch is Schorfheide-Chorin. Level 1 is the description of the codes assigned in data creation (further documentation is available at https://sg.geodatenzentrum.de/public/gdz/dokumentation/eng/lbm-de2018_eng.pdf). Level 2 shows the broader categories we grouped Level 1 values into for modeling purposes.**

| Code | Level 1 | Alb | Hai | Sch | Level 2 | Alb | Hai | Sch |
|---|---|---|---|---|---|---|---|---|
| B211 | Arable land | 17.2 | 52.0 | 23.6 | Cropland | 17.2 | 52.1 | 23.6 |
| B222 | Orchards and soft fruit | < 0.1 | 0.1 | < 0.1 | Cropland | | | |
| B231 | Homogeneous grassland | 25.4 | 12.1 | 13.3 | Grassland | 35.8 | 17.6 | 17.1 |
| B233 | Grassland with trees (< 50 %) | 6.0 | 4.1 | 2.8 | Grassland | | | |
| B321 | Inhomogeneous grassland | 4.5 | 1.4 | 0.9 | Grassland | | | |
| B310 | Reafforestation, young trees | 0.8 | 0.5 | 0.7 | Forest | 44.0 | 26.3 | 50.9 |
| B311 | Deciduous trees | 29.9 | 21.3 | 15.9 | Forest | | | |
| B312 | Coniferous trees | 8.7 | 2.5 | 28.0 | Forest | | | |
| B313 | Deciduous and coniferous trees | 3.6 | 1.2 | 5.2 | Forest | | | |
| B322 | Dwarf shrubs (heath) | 0.1 | < 0.1 | < 0.1 | Forest | | | |
| B324 | Bushes, shrubs | 0.9 | 0.9 | 1.2 | Forest | | | |
| B110 | Houses | 2.2 | 2.6 | 0.7 | Unvegetated | 2.9 | 4.0 | 8.5 |
| B121 | Facilities | 0.3 | 0.2 | 0.1 | Unvegetated | | | |
| B122 | Sealed areas without buildings | 0.1 | 0.1 | 0.1 | Unvegetated | | | |
| B242 | Mixed areas (regular structure) | 0.1 | 0.4 | 0.1 | Unvegetated | | | |
| B330 | Sand, stones, soil | 0.2 | 0.1 | 0.2 | Unvegetated | | | |
| B332 | Rock | 0 | 0.2 | 0 | Unvegetated | | | |
| B411 | Swamp | < 0.1 | < 0.1 | 1.0 | Unvegetated | | | |
| B412 | Bog | 0 | < 0.1 | < 0.1 | Unvegetated | | | |
| B413 | Swamp with bushes/trees < 50 % | 0 | < 0.1 | 0.4 | Unvegetated | | | |
| B414 | Bog with bushes/trees < 50 % | 0 | 0 | < 0.1 | Unvegetated | | | |
| B511 | Water courses | 0.1 | 0.1 | 0.3 | Unvegetated | | | |
| B512 | Water bodies | 0 | 0.2 | 5.7 | Unvegetated | | | |

**Table S5. Response scales of primary NCP-providing taxa, from (3).**

| NCP | Main trophic group | Response scale (m) |
|---|---|---|
| Acoustic diversity | Avian herbivores | 2000 |
| Birdwatching potential | Avian herbivores | 2000 |
| Caterpillar predation | Arthropod predators | 500 |
| Natural enemy abundance | Arthropod predators | 500 |
| Pollination | Insect pollinators | 2000 |
| Herbivory | Insect herbivores | 2000 |
| Pathogen infection | Fungal pathogens | 2000 |
| Seed predation | Avian herbivores | 2000 |

**Table S6. Details of the sampling methods for each NCP we tested. The number of plots represents the number of plots with non-missing data out of 150 grassland plots. The number of subplots with non-missing data out of 750 is presented in parentheses for caterpillar predation and seed predation, which were modeled at the subplot level.**

| NCP | Indicator | Year | Plots (subplots) | Data owners | Dataset ID | Sampling methods |
|---|---|---|---|---|---|---|
| Acoustic diversity | Distribution of acoustic energy among frequency bands | 2016 | 114 | S. Müller, M. Scherer-Lorenzen | 27568 (13), 27569 (14), 27570 (15) | Sounds were recorded 1 min every 10 min each day in April and May 2016, from 7 am to 7 pm, using an autonomous recording system (Soundscape Explorer T, Lunilettronics) placed at 2 m height in the center of the grassland plot. The acoustic diversity was calculated across the frequency range of 0-24 kHz using 1 kHz steps and a decibel threshold of -50. |
| Birdwatching potential | Bird species richness | Sum between 2008 and 2012 | 150 | K. Jung, S. Renner, M. Tschapka, S. Böhm | 21447 (16) | Birds were surveyed during the breeding season (March–June) by standardized audio-visual point-counts between 2008 and 2012. We used fixed-radius point counts and recorded all individuals, seen or heard during a five-minute count during the morning chorus (sunrise–11:00 h) were registered. In exceptional cases, observations were made during the evening chorus (last 3 h before sunset). Each plot was visited five times each year. |
| Caterpillar predation | Probability of dummy caterpillar being predated by birds after 48 h | 2017 | 134 (668) | N. Blüthgen, N. Simons, W. Weisser | 25806 (17) | On each plot, 5 subplots were selected at three plot corners and the middle of the two plot sides in between the corners. At each subplot, 10 dummy caterpillars were attached to the ground with insect needles. After 48 h, dummy caterpillars were retrieved and pinned unto styrofoam inside cardboard boxes. In the lab, each dummy caterpillar was inspected under a Stereo microscope and the number and type of predation marks identified. Predation marks were assigned to arthropods, birds, small mammals and gastropods. |

Continued on next page

**Table S6 – continued from previous page**

| NCP | Indicator | Year | Plots (sub-plots) | Data owners | Dataset ID | Sampling methods |
|---|---|---|---|---|---|---|
| Natural enemy abundance | Number of parasitoid predating pest insects recorded in trap-nesting wasps | 2008 | 83 | J. Steckel, C. Westphal, I. Steffan-Dewenter | 13350, 13347, 13349 (18) | Four wooden poles were placed 4 m apart on each plot and two trap nests were mounted 1.5 m high on each pole. Trap nests were constructed using PVC tubes 10.5 cm in diameter, filled with reed internodes of *Phragmites australis*. To sample the entire community of cavity-nesting species, we used reed of internodes differing in diameter (0.2–1.2 cm). Trap nests were installed between the middle of April and the middle of May 2008 and were collected at the end of September and beginning of October 2008. The traps were stored until hatching and the wasps emerging were counted and identified to species. Here we include only those wasps feeding on pest insects. This was the total number of wasp individuals belonging to the families Crabonidae (excluding *Trypoxylon* species, which feed on spiders) and Vespidae. |
| Pollination | Total abundance of flower visitors | 2008 | 119 | C. Weiner , M. Werner, N. Blüthgen | 15086 (18) | On a transect of 200×3 m along the plot edge, all individual flower visitors were recorded and identified during three transect walks (total 6 h) on a single day between April and August 2008. The total number of individuals of the orders Diptera, Hymenoptera, Lepidoptera and Coleoptera (excluding Nitidulidae) defined the total abundance used here. |
| Herbivory | Total proportion of leaf area damaged by herbivores | 2017 and 2018 | 147 | F. Neff, M. Goßner | 24086 (18) | Based on vegetation records from the previous year, we collected leaf material of the 10 most abundant plant species at the margins of each 50 × 50 m plot to reduce impact on other experiments in May 2017 or 2018. Plant material was collected before the first mowing event. For each plant, we visually estimated the area damaged by invertebrate herbivores on 12 to 200 leaves (depending on leaf size) and measured total leaf area using a leaf area meter. The deduced herbivory rates (% damaged area) per plant species were then summarized to community-level herbivory rates based on the respective plant cover values in vegetation records of the sampling year (2017 or 2018). |

Continued on next page

**Table S6 – continued from previous page**

| NCP | Indicator | Year | Plots (sub-plots) | Data owners | Dataset ID | Sampling methods |
|---|---|---|---|---|---|---|
| Pathogen infection | Total cover of foliar fungal pathogens | 2011 | 142 | S. Blaser, D. Prati, M. Fischer | 24247 (18) | On four transects of $25 \times 1$ m per plot all plant species were scanned for pathogens infection, including rust, powdery mildew, downy mildew and smut fungi between May and June 2011. The percentage of infected plants was multiplied with the severity per pathogen species (divided by 1000 to get a number between 0 and 1). The infection of all pathogens per plant species was combined, because one plant species can be infected by various pathogens at the same time. The infection severity per plant species was multiplied with the according plant species cover on each plot separately. |
| Seed predation | Probability of sunflower seed being removed after 48 h | 2017 | 130 (624) | N. Blüthgen, K. Wehner | 24966 (19) | Twenty-five nonviable intact sunflower seeds were placed on plastic trays (reduce loss by wind). After 48 h remaining seeds were counted. Due to the effect of rain washing away seeds, data from sites with more than 30 mL precipitation within the 48 h were excluded (18 sites). |

**Table S7. Structure of full controls model for each NCP (model 4 or model 2 in *SI Appendix* Table S2 for NCP respectively with and without controls for other NCP). Here, $i$ is plot, $j$ is subplot, $\text{TWI}_i$ is topographic wetness index. Caterpillar predation and seed predation were measured and therefore modeled at the subplot level.**

| NCP | Model likelihood | Model equation |
|---|---|---|
| Acoustic diversity | $y_i \sim Beta(\mu, \phi)$ | $\text{logit}(\mu_i) = \beta_{0,\text{region}} + \beta_{1,\text{region}}\text{TWI}_i + \beta_{2,\text{region}}\text{Birdwatching potential}_i$ |
| Birdwatching potential | $y_i \sim Poisson(\lambda_i)$ | $\log(\lambda_i) = \beta_{0,\text{region}} + \beta_{1,\text{region}}\text{TWI}_i$ |
| Caterpillar predation | $y_{i,j} \sim Binomial(\text{N}, \theta_{i,j})$ | $\text{logit}(\theta_{i,j}) = \beta_{0,\text{region}} + \beta_{1,\text{region}}\text{TWI}_i + \beta_{2,\text{region}}\text{Birdwatching potential}_i$ |
| Natural enemy abundance | $y_i \sim Poisson(\lambda_i)$ | $\log(\lambda_i) = \beta_{0,\text{region}} + \beta_{1,\text{region}}\text{TWI}_i$ |
| Pollination | $y_i \sim Poisson(\lambda_i)$ | $\log(\lambda_i) = \beta_{0,\text{region}} + \beta_{1,\text{region}}\text{TWI}_i$ |
| Herbivory | $y_i \sim Beta(\mu, \phi)$ | $\text{logit}(\mu_i) = \beta_{0,\text{region}} + \beta_{1,\text{region}}\text{TWI}_i$ |
| Pathogen infection | $y_i \sim Beta(\mu, \phi)$ | $\text{logit}(\mu_i) = \beta_{0,\text{region}} + \beta_{1,\text{region}}\text{TWI}_i + \beta_{2,\text{region}}\text{Herbivory}_i$ |
| Seed predation | $y_{i,j} \sim Binomial(\text{N}, \theta_{i,j})$ | $\text{logit}(\theta_{i,j}) = \beta_{0,\text{region}} + \beta_{1,\text{region}}\text{TWI}_i + \beta_{2,\text{region}}\text{Birdwatching potential}_i$ |

### References


1. G Peralta, et al., Scale-dependent effects of landscape structure on pollinator traits, species interactions and pollination success. *Ecography* **2023**, e06453 (2023).
2. SJ Hegland, Floral neighbourhood effects on pollination success in red clover are scale-dependent. *Funct. Ecol.* **28**, 561–568 (2014).
3. G Le Provost, et al., Contrasting responses of above-and belowground diversity to multiple components of land-use intensity. *Nat. Commun.* **12**, 1–13 (2021).
4. S Müller, et al., Temporal dynamics of acoustic diversity in managed forests. *Front. Ecol. Evol.* **12**, 1392882 (2024).
5. P Anttonen, et al., Predation pressure by arthropods, birds, and rodents is interactively shaped by tree species richness, vegetation structure, and season. *Front. Ecol. Evol.* **11**, 1199670 (2023).
6. JG Lundgren, JT Shaw, ER Zaborski, CE Eastman, The influence of organic transition systems on beneficial ground-dwelling arthropods and predation of insects and weed seeds. *Renew. Agric. Food Syst.* **21**, 227–237 (2006).
7. D Vujanović, et al., Forest and grassland habitats support pollinator diversity more than wildflowers and sunflower monoculture. *Ecol. Entomol.* **48**, 421–432 (2023).
8. E González, DA Landis, M Knapp, G Valladares, Forest cover and proximity decrease herbivory and increase crop yield via enhanced natural enemies in soybean fields. *J. Appl. Ecol.* **57**, 2296–2306 (2020).
9. H Susi, AL Laine, Agricultural land use disrupts biodiversity mediation of virus infections in wild plant populations. *New Phytol.* **230**, 2447–2458 (2021).
10. MM Gossner, L Beenken, K Arend, D Begerow, D Peršoh, Insect herbivory facilitates the establishment of an invasive plant pathogen. *ISME Commun.* **1**, 6 (2021).
11. K Wehner, L Schäfer, N Blüthgen, K Mody, Seed type, habitat and time of day influence post-dispersal seed removal in temperate ecosystems. *PeerJ* **8**, e8769 (2020).
12. N Breitbach, I Laube, I Steffan-Dewenter, K Böhning-Gaese, Bird diversity and seed dispersal along a human land-use gradient: high seed removal in structurally simple farmland. *Oecologia* **162**, 965–976 (2010).
13. M Scherer-Lorenzen, S Mueller, Acoustic diversity index based on environmental sound recordings on all forest eps, hai, 2016. biodiversity exploratories information system. https://www.bexis.uni-jena.de/ddm/data/Showdata/27568 (2023).
14. M Scherer-Lorenzen, Acoustic diversity index based on environmental sound recordings on all forest eps, alb, 2016. biodiversity exploratories information system. https://www.bexis.uni-jena.de/ddm/data/Showdata/27569 (2023).
15. M Scherer-Lorenzen, Acoustic diversity index based on environmental sound recordings on all forest eps, sch, 2016. biodiversity exploratories information system. https://www.bexis.uni-jena.de/ddm/data/Showdata/27570 (2023).
16. M Tschapka, S Renner, K Jung, Bird survey data 2009, all 300 ep. biodiversity exploratories information system. https://www.bexis.uni-jena.de/ddm/data/Showdata/21447 (2020).
17. N Simons, K Wehner, N Blüthgen, W Weisser, Predation marks on dummy caterpillars measured on all eps in 2017. biodiversity exploratories information system. https://www.bexis.uni-jena.de/ddm/data/Showdata/25806 (2021).
18. NV Schenk, C Penone, E Allan, M Fischer, Assembled ecosystem measures from grassland eps (2008-2018) for multi-functionality synthesis - june 2020. biodiversity exploratories information system. https://www.bexis.uni-jena.de/ddm/data/Showdata/31621 (2023).
19. N Blüthgen, K Wehner, Arthropod mediated processes (dung and seed depletion) measured on all eps in 2017. biodiversity exploratories information system. https://www.bexis.uni-jena.de/ddm/data/Showdata/24966 (2021).
